# Parallel evolution of a splicing program controlling neuronal excitability in flies and mammals

**DOI:** 10.1101/2021.02.24.432780

**Authors:** Antonio Torres-Méndez, Sinziana Pop, Sophie Bonnal, Isabel Almudi, Alida Avola, Ruairí J V Roberts, Chiara Paolantoni, Ana Alcaina, Ane Martín-Anduaga, Irmgard U Haussmann, Violeta Morin, Fernando Casares, Matthias Soller, Sebastian Kadener, Jean-Yves Roignant, Lucia Prieto-Godino, Manuel Irimia

**Affiliations:** Centre for Genomic Regulation, Barcelona Institute of Science and Technology (BIST), Barcelona 08003, Spain; The Francis Crick Institute, London, UK; Centro Andaluz de Biología del Desarrollo (CABD), Seville, Spain; Center for Integrative Genomics, Génopode Building, Faculty of Biology and Medicine, University of Lausanne, CH-1015, Lausanne, Switzerland; Biology Department, Brandeis University, Waltham, United States; Department of Life Science, School of Health Sciences, Birmingham City University, Birmingham, B5 3TN, UK; Institute of Molecular Biology (IMB), Mainz, Germany; School of Biosciences, College of Life and Environmental Sciences, University of Birmingham, Edgbaston, Birmingham, B15 2TT, UK; Birmingham Centre for Genome Biology, University of Birmingham, Edgbaston, Birmingham, B15 2TT, UK; Universitat Pompeu Fabra (UPF), Barcelona 08003, Spain; ICREA, Barcelona, Spain

**Keywords:** evolution, splicing, neuron, microexons, Drosophila, mouse, ion channels

## Abstract

Neurons draw on alternative splicing for their increased transcriptomic complexity throughout animal phylogeny. To delve into the mechanisms controlling the assembly and evolution of this regulatory layer, we characterized the neuronal microexon program in *Drosophila* and compared it with that of mammals. We found that in *Drosophila*, this splicing program is restricted to neurons by the post-transcriptional processing of the *enhancer of microexons* (eMIC) domain in *Srrm234* by Elav and Fne. eMIC deficiency or misexpression leads to widespread neurological alterations largely emerging from impaired neuronal activity, as revealed by a combination of neuronal imaging experiments and cell-type-specific rescues. These defects are associated with the genome-wide skipping of short neural exons, which are strongly enriched in ion channels. Remarkably, we found no overlap of eMIC-regulated exons between flies and mice, illustrating how ancient post-transcriptional programs can evolve independently in different phyla to impact distinct cellular modules while maintaining cell-type specificity.

## Introduction

Protein-coding genes in metazoans undergo multiple mRNA processing steps before they are ready for translation. One pivotal step is the removal of introns, mediated by the interaction of the splicing machinery and other related proteins with the pre-mRNA ^1^. Splice site selection is not deterministic and indeed several mRNA products can be produced from the same gene in a process known as alternative splicing (AS). AS can greatly expand the coding capacity of metazoan genomes ^2^, with striking examples including the *Down syndrome adhesion molecule 1* (Dscam1) from *Drosophila melanogaster*, which can generate over 35,000 AS isoforms from a single gene ^3^.

In evolutionary terms, AS can serve similar functions as gene duplication since it allows for the exploration of new coding capabilities without affecting pre-existing gene functionality ^4,5^. In metazoans, neural tissues have particularly exploited the potential brought by AS and present the highest number of tissue-enriched exons ^6–11^. These neural enriched isoforms have been implicated in key aspects of neuronal biology including neurogenesis, axon guidance and growth, synapse formation and synaptic plasticity ^12–14^. Neural splicing programs are coordinated by the action of RNA binding proteins (RBPs) that are predominantly expressed in this tissue and can modulate hundreds of splicing decisions genome-wide ^15^. The importance of these splicing choices in the brain is underscored by the widespread association between splicing alterations and neurological disorders such as in autism spectrum disorder, spinal muscular atrophy, amyotrophic lateral sclerosis, Huntington’s disease or intellectual disability, among others ^16–19^.

Among these programs, transcriptomic analyses across vertebrate tissues and human brain samples uncovered a highly conserved set of very short neural-enriched exons: microexons. These are down-regulated in some autistic patients ^20^, and their mis-regulation in mouse models leads to a wide range of neurological phenotypes ^21–23^. Splicing of neural microexons is regulated by the combinatorial action of several splicing factors. The Serine/Arginine Repetitive Matrix 4 protein SRRM4 and its paralogue SRRM3 are the master regulators of microexon splicing, being sufficient to promote inclusion of ~90% of neural microexons when ectopically expressed in non-neural cells ^20,24^. Many neural microexons are also repressed by PTBP1 in non-neural samples ^25,26^, thereby reinforcing their switch-like profile across tissues. A recent high-throughput study searching for microexon regulators identified two additional factors, RNPS1 and SRSF11, which cooperate with SRRM4 to assemble an exon definition complex that facilitates microexon splicing ^27^. Expanding microexon profiling beyond vertebrates revealed that neural microexons originated in bilaterian ancestors in association with the appearance of a novel domain in the ancestral *Srrm234* gene that is necessary and sufficient for neural microexon splicing: the *enhancer of microexons* or ‘eMIC’ domain ^24^. Neural expression of the eMIC domain is regulated transcriptionally through the expression of *Srrm3* and *Srrm4* in vertebrates, both containing the eMIC domain, but through post-transcriptional processing of *Srrm234* in non-vertebrates ^24^.

Here, we address the regulation, functional impact and evolution of the neural microexon program in a non-vertebrate. For this, we generated *D. melanogaster* flies with eMIC loss-of-function, as well as transgenic lines for cell-type-specific expression of different variants of the *Srrm234* gene. eMIC-null flies display an array of neurological defects, including alterations in locomotion, ageing, sleep, metabolism and bang-sensitivity. Expression of transgenic *Srrm234* variants in different cell types underscores the relevance of spatially and quantitatively regulating eMIC activity in *Drosophila*. Importantly, lack of eMIC activity results in genome-wide down-regulation of AS exons, affecting up to one third of all neural exons in *D. melanogaster*. By profiling AS in over 700 RNA-seq samples we generated a catalogue of all tissue and cell-type specific exons, with a focus on eMIC-dependent exons. We also characterized the *cis*-regulatory code associated with eMIC-dependent splicing in *Drosophila*, highlighting differences and similarities with mammals. Strikingly, despite the remarkable cell type-specific conservation, we only found four exons in equivalent positions between the fly and mammalian eMIC splicing programs, indicating that both programs evolved largely independently from an ancestral neuronal-specific program.

## Results

### Regulated 3′ end processing of *Srrm234* ensures strict eMIC neural expression

The *Srrm234* gene in *Drosophila* can produce disparate protein isoforms based on the post-transcriptional processing at its 3′ end (Figure 1A and Supp. Figure 1). Alternative last exon selection at this locus depends on a combination of AS and alternative poly-adenylation (APA) events (Figure 1A). The proximal non-eMIC-encoding exon can be expressed either as a terminal exon making use of its own poly-A site (pA_1_ site in Figure 1A, isoform A) or as a “poison” exon for the eMIC-expressing isoform when the distal poly-A site is used (pA_2_ site in Figure 1A, isoform G). Translation of the distal exon encoding the eMIC domain requires both distal poly-A usage and skipping of the proximal exon (C and F isoforms). This particular genetic architecture encoding the eMIC domain as an alternative last exon of the *Srrm234* gene is conserved at least within holometabolous insects (Figure 1B).

**Figure 1.**
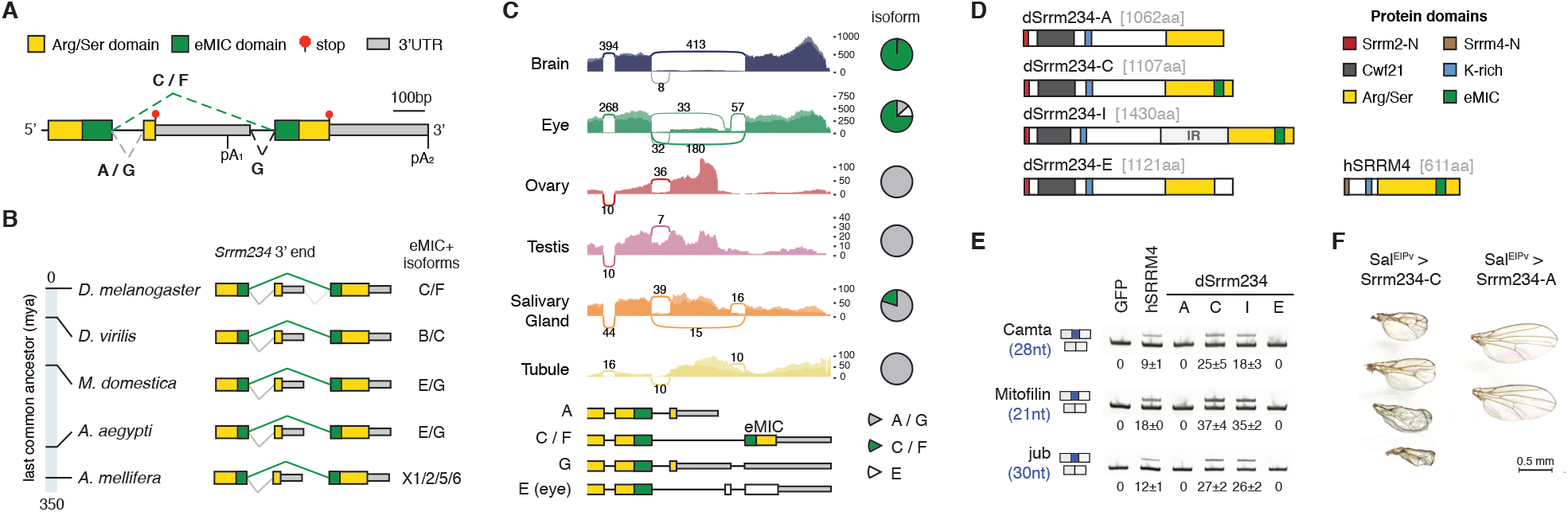
Post-transcriptional regulation of the neural expression of the eMIC domain in *D. melanogaster*. **A.** Genomic region encompassing the 3′-most terminal exons of the *Srrm234* gene and the corresponding protein domains encoded therein. Arg/Ser: Arginine/Serine-rich, eMIC: enhancer of microexons, 3′UTR: 3′ untranslated region. Putative AS and poly-adenylation (pA) sites are indicated together with their associated reference transcripts (A, C, F, G). **B.** Genomic architecture of the 3′ end of the *Srrm234* locus in different insect species. Reference isoforms encoding the eMIC domain are indicated. mya: million years ago. **C.** Sashimi plots of RNA-seq data from different tissues at *Srrm234* 3′ end region. Average numbers of reads spanning each splice junction between male and female samples are indicated. Y-axis represents absolute number of mapping reads (without normalization for library size). Bottom: main transcript isoforms annotated for the *Srrm234* gene (FlyBase annotation). Pie charts depict isoform usage quantified based on junction reads only. Data from FlyAtlas 2 ^28^. **D.** Protein domains of *Drosophila* (d) Srrm234 isoforms and human (h) SRRM4. IR: intron retention, aa: amino acids, K: lysine, Srrm2/4-N: conserved regions at Srrm2/4 N-termini. **E.** RT-PCR assays for alternatively spliced exons in SL2 cells overexpressing different Srrm234 isoforms. **F.** Representative pictures of fly wings overexpressing either UAS-Srrm234-C or -A under the control of *spalt* (Sal^E|PV^-GAL4) driver line, active in the centre of the wing blade ^29^.

We analysed isoform usage at this region using tissue-specific RNA-seq data from FlyAtlas 2 ^28^ and found that eMIC expression (isoforms C/F) is strongly biased towards neural tissues (brain and eyes), whereas other tissues mainly express *Srrm234* isoforms with no eMIC (A/G) (Figure 1C and Supp. Figure 1). This analysis also revealed a fourth splice variant that is only expressed in the eye (Figure 1C). We cloned representative *Srrm234* isoforms to test their splicing activity on neural microexons by heterologous expression in *Drosophila* SL2 cells (Figure 1D). We chose four isoforms for this experiment: the two reference isoforms A and C, which only differ at the C-term where the full eMIC is encoded in isoform C but not in A; a variant of isoform C containing a protein-coding neural-retained intron (IR) as in reference isoform F, which we termed isoform I; and the newly identified eye-specific isoform, that we named isoform E. As a control, we used the human ortholog SRRM4 (hSRRM4), which we have previously shown to promote inclusion of short endogenous neural exons in this system ^24^. Consistent with previous studies, only the proteins harbouring the complete eMIC domain were able to promote inclusion of short neural exons (Figure 1E and Supp. Figure 2A).

To investigate the functional relevance of the restriction of the eMIC to neural tissues, we ectopically expressed the eMIC domain in a non-neural tissue (the wing) by generating transgenic flies with different *Srrm234* isoforms under the control of the GAL4-specific UAS enhancer (Figure 1F and Supp. Figure 2B). Expression of isoform C, but not A, in the entire wing pouch under a *nubbin* (*nub*) driver line prevented formation of adult wings and severely affected haltere morphology (Supp. Figure 2B). Expression in the centre of the wing blade only, under a *spalt* (*Sal^E|PV^*) driver ^29^, generated bubbles, shortening and blister phenotypes (Figure 1F). These results highlight the detrimental effects of eMIC expression outside neuronal tissues, and the latter suggests additional non-cell autonomous effects, as defects spread beyond the delimited area of *Sal^E|PV^* expression. Together, these results show that *Srrm234* last exon selection, and hence eMIC domain expression, needs to be tightly regulated to restrict its activity to the neural system.

### Altered eMIC expression levels results in widespread neurological defects in *Drosophila*

To characterize the neural microexon splicing program in *Drosophila*, we generated eMIC-specific knockout flies (*Srrm234*^eMIC-^; hereafter eMIC-) via CRISPR-Cas9 targeting the 3′ region of *Srrm234* with two gRNAs and replacing it with an integration cassette (Figure 2A, Supp. Figure 2C and Methods). Only ~15% of eMIC-flies reach pupal stage. Furthermore, surviving eMIC-adult flies are smaller than controls, with a 20% reduction in body weight at hatching, and have reduced lifespan (Figure 2B-D). These size and weight reductions are correlated with reduced levels of neuronally secreted insulin-like peptides: Ilp2, Ilp3 and Ilp5 ^30^ (Supp. Figure 2D). As expected, RT-PCR assays on fly heads showed that eMIC insufficiency leads to skipping of known short neural-enriched exons (Supp. Figure 2E).

**Figure 2.**
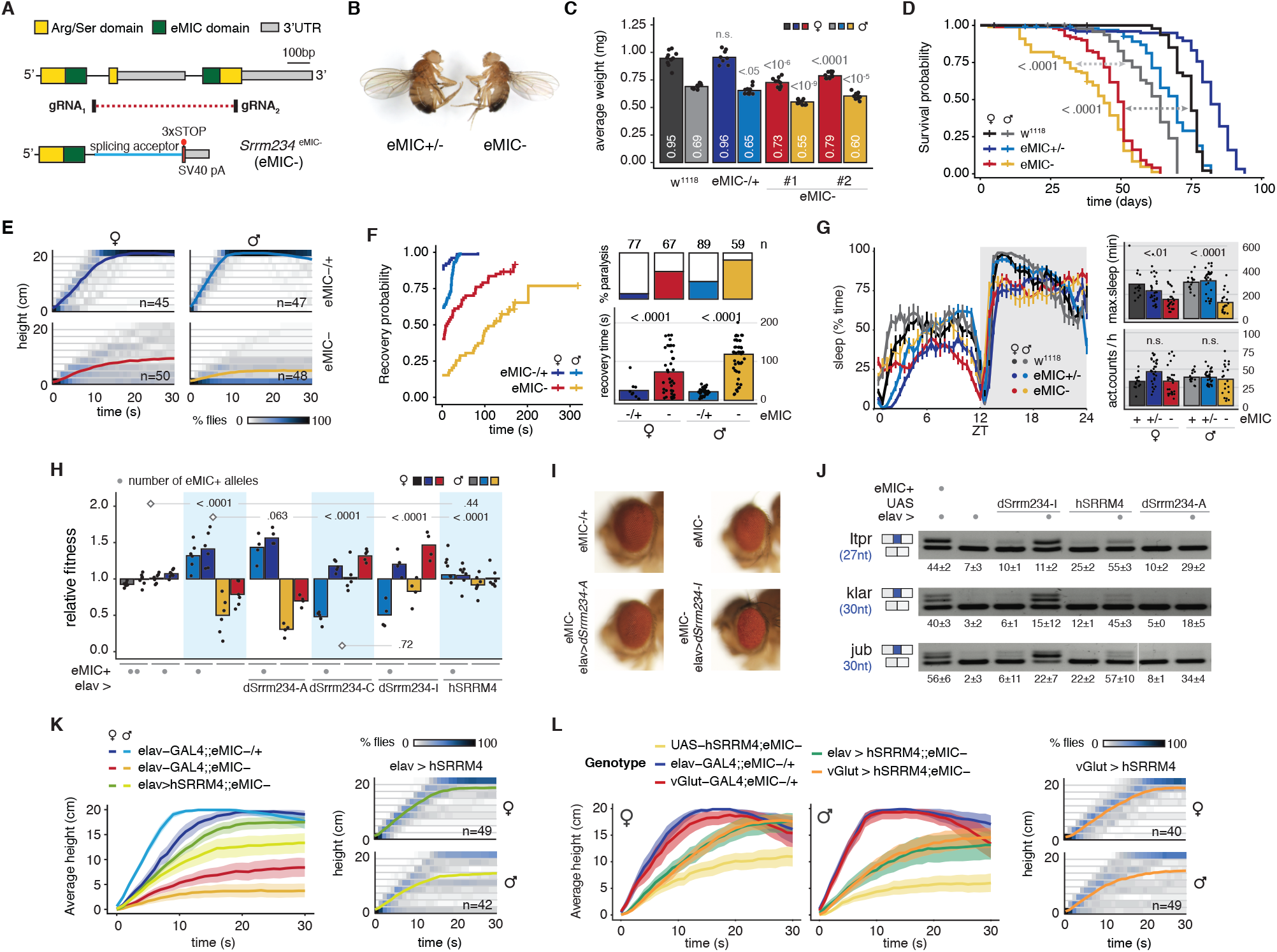
Physiological alterations associated with eMIC domain loss or misexpression. **A.** CRISPR strategy to generate mutant flies with eMIC-specific deletion at the *Srrm234* locus. The deleted part of the ‘enhancer of microexons’ (eMIC) protein domain is encoded in the alternative last exon and is essential for its function. The dotted red line spans the deleted genomic region, which was replaced by an integration cassette (Supp. Figure 2C). Arg/Ser: Arginine/Serine-rich, eMIC: enhancer of microexons, 3′UTR: 3′ untranslated region, gRNA: guide RNA, pA: poly-adenylation site. **B.** Left: eMIC+/− male 1 day-old fly from the cross with w^1118^. Right: eMIC-male. **C.** Average weight of w^1118^ controls, eMIC-/+ heterozygous flies and two independent eMIC-clones (#1, #2) from the CRISPR targeting, less than 24 hours after hatching. White numbers indicate mean values. P-values from two-sided t-tests comparing with w^1118^ controls, n.s.: non-significant or p > 0.05. **D.** Longevity assay, n=100 flies per sex and genotype. P-value from log-rank tests of eMIC-flies compared to controls for each sex. **E.** Kymographs displaying *Drosophila* negative geotaxis behaviour. Coloured lines indicate average height at each timepoint. **F.** Sensitivity of adult flies to mechanical stimulation (10s vortex). Left: probability of recovering from mechanical-induced paralysis over time. Top right: number of flies tested and percentage of them that are sensitive to mechanical stress. Bottom right: time spent in recovering from paralysis until the fly is in upright position. P-values from Mann-Whitney U-tests. **G.** Sleep patterns of 3 day-old flies in 12h light – 12h dark cycles. Left: average time flies spend sleeping (inactive for >= 5min) at different times of the light:dark cycle. ZT: Zeitbeger Time (switch from light to dark conditions), vertical lines: standard error of the mean. Top right: maximum sleep episode during the night. Bottom right: total number of activity counts per hour during the night. P-value from Mann-Whitney U-tests comparing eMIC-with control flies, n.s.: p > 0.05. **H.** Relative fitness of flies with varying number of eMIC+ alleles in the F1 generation. See Supplementary Figure 2F for a detailed mating scheme. P-values from χ^2^-tests on the observed frequencies of genotypes per cross, compared with crosses marked with a diamond. Transgenic construct expression is induced pan-neuronally using an elav-GAL4 driver. **I.** Representative pictures of eyes from young flies expressing *dSrrm234* isoforms under an elav-GAL4 driver in the eMIC-background. **J.** RT-PCR assays of short neural exons from female fly heads. Numbers indicate mean and standard deviation values from three replicates. **K-L.** Kymographs displaying the negative geotaxis behaviour of *Drosophila* eMIC-flies upon expression of UAS-*hSRRM4* pan-neuronally using an elav-GAL4 driver line (K-L) or in glutamatergic neurons (including motoneurons) under a vGlut-GAL4 line (L). Coloured lines indicate average height at each timepoint and ribbons, 95% non-parametric bootstrap confidence intervals.

To investigate the functional role of the AS program regulated by eMIC activity, we ran a battery of behavioural assays on eMIC deficient flies. Both male and female eMIC-flies have tremors and defects in self-righting – a complex motor sequence that allows animals to adopt a locomotion position if turned upside down (Video S1). Performance of eMIC-flies in the negative geotaxis assay is very poor in both sexes, but more pronounced in males (Figure 2E). eMIC-flies are also bang sensitive, i.e. they undergo seizures after mechanical stress, with a recovery time similar to classical bang sensitive mutants ^31^ (Figure 2F). We monitored daily activity patterns and found that, despite these alterations, overall activity was similar in control and eMIC-null flies (Figure 2G). However, mutant flies sleep less and have more fragmented sleep, a phenotype that becomes more pronounced in older flies (Figure 2G and Supp. Figure 2F).

Using the GAL4-UAS system, we performed rescue experiments by expressing *Srrm234* proteins pan-neuronally (elav-GAL4) in the eMIC-background. We first calculated the relative fitness of each of the genotypes per cross, defined as the proportion of emerging adults of each genotype over the expected Mendelian proportions (Figure 2H and Supp. Figure 2G). eMIC-flies expressing *dSrrm234-A* in neurons showed the same low fitness of eMIC-flies relative to their eMIC-/+ controls, demonstrating that the A isoform lacking the eMIC domain cannot rescue eMIC mutant phenotypes (Figure 2H). Neuronal expression of *Srrm234* isoforms C and I could overcome fitness defects in eMIC-flies. However, unexpectedly, the relative fitness of control (eMIC +/−) flies overexpressing these protein forms was significantly reduced, indicating a deleterious effect associated with excessive levels of eMIC activity, particularly in males (Figure 2H and Supp. Figure 2G). Moreover, despite the increased relative fitness, *dSrrm234-C* and *-I* pan-neuronal overexpression also caused several neurological alterations, including rough eye patterning (Figure 2I), problems with wing expansion and wing and leg positioning (Supp. Figure 2H), and severe locomotion defects (Video S2). Consistent with these observations, several short neural exons had increased inclusion levels in heads of eMIC-flies with *dSrrm234-I* pan-neuronal rescue compared to controls (Figure 2J and Supp. Figure 2I). However, eMIC expression levels in the head were not higher than controls (Supp. Figure J), suggesting that the deleterious effect may come from abnormal expression in specific neuronal populations.

Beyond these, pan-neuronal expression of the human ortholog (*hSRRM4*) showed different phenotypic rescues. Specifically, pan-neuronal rescue with *hSRRM4* overcomes the relative fitness defects of eMIC-flies, despite affecting overall viability (Figure 2H and Supp. Figure 2G). Moreover, these flies have significantly improved performance on negative geotaxis assays compared to eMIC-in both sexes (Figure 2K). These rescues are associated with lower exon inclusion levels that are presumably more physiologically adequate than those induced by the misexpression of the individual fly *Srrm234* isoforms (Figure 2J and Supp. Figure 2I). Altogether, these experiments show that, beyond cell-type restriction, eMIC activity needs to be tightly regulated within neurons for their correct function.

### The eMIC AS program regulates neuronal excitability

With the aim of identifying the relative share of network vs. cell-autonomous effects on eMIC-neurological alterations, we next rescued eMIC expression in eMIC-flies only in glutamatergic neurons (which includes motoneurons and a restricted number of interneurons) by expressing *hSRRM4* under the regulation of vGlut-GAL4. Surprisingly, rescue in glutamatergic neurons alone restored climbing performance to the same extent as pan-neuronal rescue (Figure 2L), indicating that the contribution of eMIC insufficiency to that phenotype mainly stems from cell-autonomous alterations in this neuronal population.

To further investigate the mode of action of the eMIC-regulated AS program, we then focused on the *Drosophila* larvae. We first assessed free crawling behaviour in third instar larvae, and found that the neurological-associated phenotypes were also present at this stage: eMIC-larvae crawl more slowly, perform more turns with less straight paths, and display unusual unilateral body-wall contractions that lead to C-shape behaviour (Figure 3A). Similar to adult climbing assays, these phenotypes could be partially rescued by pan-neuronal expression of *hSRRM4* (Figure 3B). Despite the abnormal crawling behaviour, examination of overall central nervous system (CNS) morphology revealed no differences between control and eMIC-larvae (Figure 3C). Moreover, single-cell RNA-seq of eMIC- and control L1 CNS at ~2X coverage (~19,000 cells per genotype) revealed no defects in the generation of major cell types (Figure 3D), which could be readily identified based on known marker genes (Supp. Figure 3). We further examined in detail motoneuron axonal terminals and the synapses between motoneurons and muscles at the neuromuscular junction (NMJ), and also found no alterations in eMIC-larvae (Figure 3E). Altogether, these results indicate that abnormal crawling behaviour is not due to major developmental or morphological alterations.

**Figure 3.**
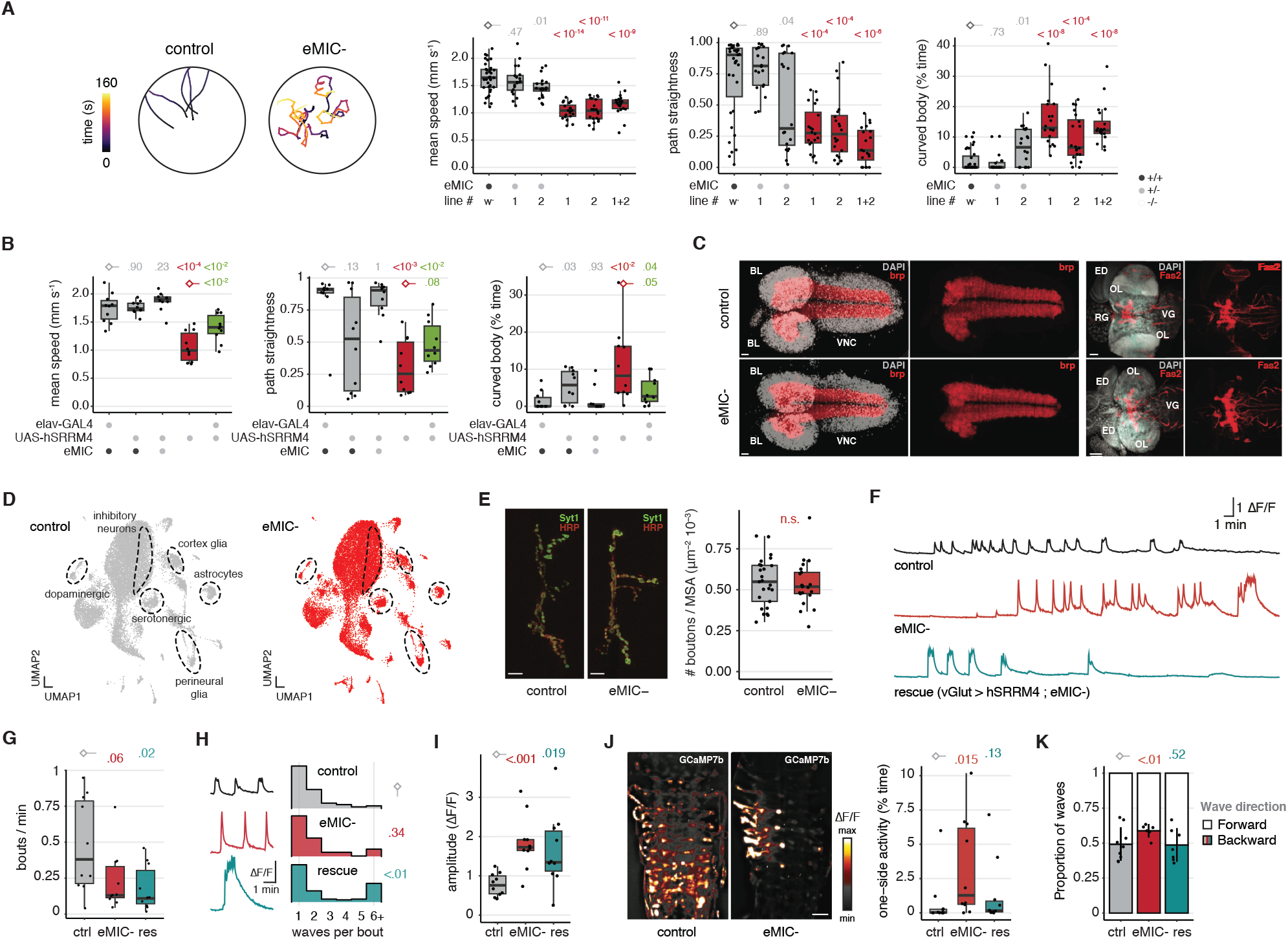
Alterations in locomotion behaviour and neuronal activity in eMIC-larvae. **A.** Left: Locomotion tracks of free crawling third instar larvae (L3). Right: boxplots for the quantifications of different parameters describing larval locomotion. #1-#2 corresponds to the F1 trans-heterozygous from crossing two independent eMIC-lines from the CRISPR-Cas9 genome editing. **B.** Quantification of L3 free crawling behaviour upon expression of human *SRRM4* (*hSRRM4*) pan-neuronally using an elav-GAL4 driver. P-values from Welch (speed) or Mann-Whitney U tests (path straightness and curved body patterns) comparing to w^1118^ controls. **C.** Confocal images of control and eMIC-L3 CNSs using antibodies against Bruchpilot (brp, nc82) and Fasciclin 2 (Fas2). BL: brain lobe, VNC: ventral nerve cord, OL: optic lobe, VG: ventral ganglia, ED: eye disc, RG: ring gland. Size bar: 20 μm and 50 μm in brp and Fas2 stainings, respectively. **D.** Single-cell RNA-sequencing data visualized using the uniform manifold approximation and projection (UMAP) algorithm after integrating control and mutant datasets. Identification of cell populations is based on the following markers: *Tdc2* (dopaminergic), *SerT* (serotonergic), *Gad1* (inhibitory neurons), *wrapper* and *hoe1* (cortex glia), *CG6126* and *Indy* (perineural glia), *alrm* and *Gat* (astrocytes). **E.** Synaptic bouton quantification in the larval neuromuscular junction for control and eMIC-flies based on immunostaining against synaptotagmin (Syt1), and anti-HRP to mark neuronal membranes. MSA: muscle surface area. Size bar: 10 μm. **F.** Example traces representing the mean activity across all segments of the ventral nerve cord (VNC) in fictive locomotion experiments using GCaMP7b calcium indicator for control, eMIC- and rescue (vGlut > *hSRRM4*; eMIC-) larvae. F: fluorescence. **G-I.** Parameters describing the activity patterns of glutamatergic neurons in the VNC during fictive locomotion experiments: number of bouts per minute (G), number of waves per bout (H) and peak amplitude (I). P-values from Welch tests (bouts/min and amplitude) or Mann-Whitney U tests (waves per bout). **J.** One-sided activity events in the VNC during fictive locomotion experiments. Left: example of one-sided activity in eMIC-larval VNC compared to symmetric activity in the control. Size bar: 25 μm. Right: quantification of the frequency of those events for each genotype. P-values from Mann-Whitney U tests. **K.** Proportion of forward and backward waves during fictive locomotion. P-values from logistic regression. res: rescue line expressing *hSRRM4* in glutamatergic neurons in the eMIC-background (vGlut > *hSRRM4*; eMIC-).

Therefore, we reasoned that motor phenotypes could be due to defects in neuronal activity. To test this hypothesis, we examined motoneuron activity in the ventral nerve cord by expressing a genetically encoded calcium indicator (UAS-GCaMP7b) in all glutamatergic neurons (vGlut-GAL4). When isolated, *Drosophila* larval CNS produces spontaneous ventral nerve cord activity patterns that recapitulate the sequence of muscle activation during locomotion, a process referred to as fictive locomotion ^32^. Neuronal activation correlates with turning and crawling, albeit it is ten times slower than during actual behavioural sequences. These activity patterns occur during activity bouts separated by non-active periods ^32,33^ (Figure 3F and Supp. Figure 4A). We found that the CNSs of eMIC-larvae generated activity bouts at slightly reduced rates compared to controls, and that this phenotype could not be rescued by expressing *hSRRM4* in glutamatergic neurons (Figure 3G). However, whereas control and eMIC-CNSs generated similar number of waves per bout, vGlut *hSRRM4* rescues partially compensated for the low number of bouts by increasing the number of waves per bouts (Figure 3H). This indicates that, while the number of bouts might be a property of the whole network, neurons with a functioning eMIC program can cell-autonomously regulate their excitability to generate a higher number of waves. Supporting the dysregulation of neuronal excitability in eMIC-larvae, we found that their calcium waves had significantly higher amplitudes than control (Figure 3I, Supp. Figure 4B and Video S3). This phenotype was partially, but not completely, rescued by the expression of *hSRRM4* (Figure 3I). Finally, we found that eMIC-CNSs generated a high number of spontaneous unilateral activity events, mirroring our behavioural experiments, where eMIC-larvae displayed unusual unilateral body-wall contractions (Figure 3J). Moreover, it also generated a higher proportion of backward waves (Figure 3K and Supp. Figure 4C). Notably, these phenotypes could be fully rescued by restoring eMIC expression with *hSRRM4* in glutamatergic neurons (Figure 3J,K). Altogether, these results point to an important role of the eMIC-regulated AS program in controlling neuronal activity.

### The eMIC domain regulates short neural exons genome-wide in *Drosophila*

To comprehensively characterise the splicing program regulated by the eMIC domain in *D. melanogaster* we sequenced eMIC- and control adult brains and larval CNSs, and quantified AS using *vast-tools* ^20,24^. Focusing on AS events within coding sequences, the predominant type of AS affected were cassette exons and the introns surrounding them (Figure 4A and Supp. Figure 5A). The vast majority of these exons showed increased skipping in the mutant samples (161 out of 173 regulated exons, Figure 4A-C), highlighting the role of the eMIC as a positive regulator of exon inclusion, as it has been described for Srrm3 and Srrm4 in mammals (Supp. Figure 5B and ^21,23,24,26^). Similar results were obtained when using brain samples from FlyAtlas 2 as independent controls, in line with a mutation-specific effect with little-to-none strain specificity (Supp. Figure 5C). Next, we classified exons into three groups based on their splicing pattern: eMIC-dependent (170 exons), eMIC-sensitive (128 exons) and non-eMIC-regulated exons (Figure 4C and Methods). These exons showed similar inclusion levels between males and females (Figure 4D), with only 18 eMIC-dependent exons having some mild sex differences in either control or eMIC-adult brains (Supp. Figure 5D). Similar to results from mouse Srrm4 targets ^24^, *Drosophila* eMIC targets are much shorter than other AS exons and that constitutive exons (Figure 4E), corroborating its ancestral role in regulating the inclusion of microexons.

**Figure 4.**
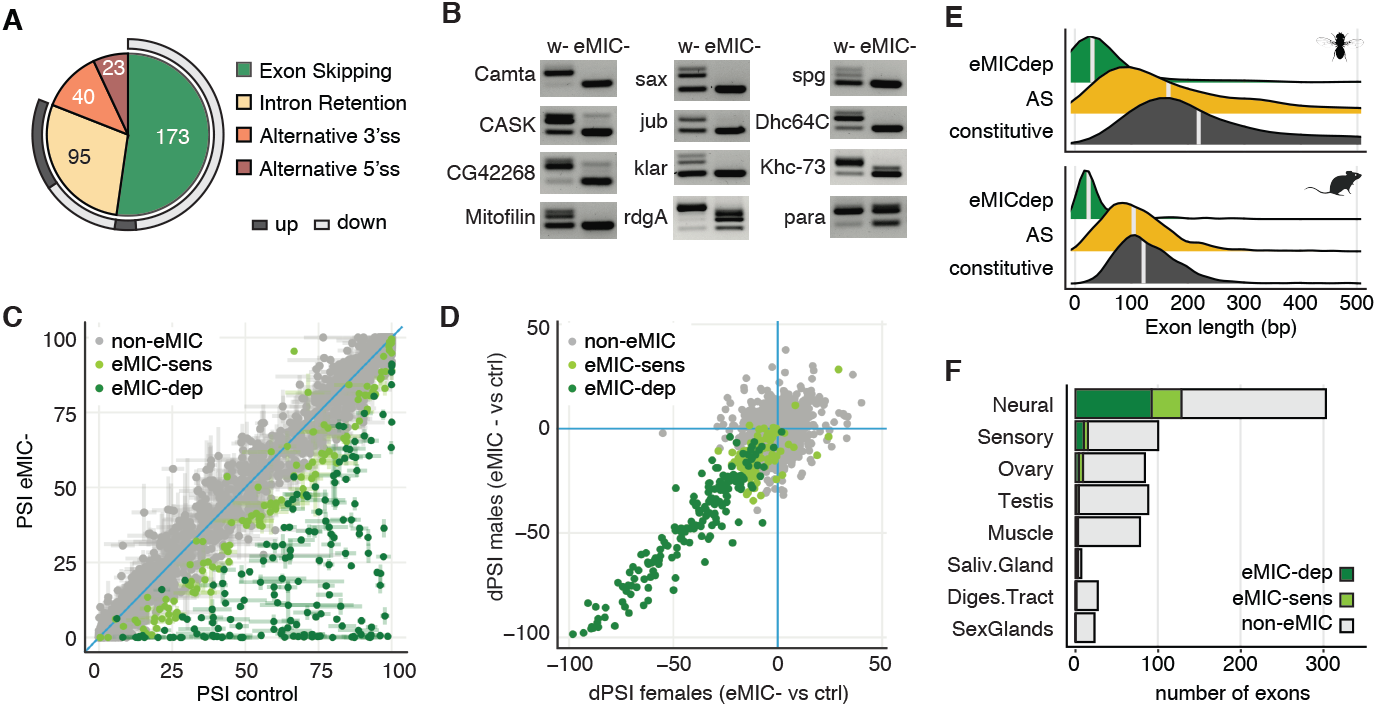
Genome-wide alternative exon inclusion regulated by the eMIC domain. **A.** Number of AS events affected by eMIC insufficiency in adult brains. ss: splice site, up or down: higher or lower inclusion/retention in eMIC-samples, respectively. **B.** RT-PCR validations of eMIC-dependent exons in control (w-) and eMIC-fly heads. **C.** Genome-wide splicing alterations for alternatively spliced (AS) exons in eMIC-adult fly brains. Error bar ends mark the PSI values from male and female samples independently. Depending on their degree of skipping upon eMIC deletion, exons are classified into three groups: eMIC-dependent (eMIC-dep), eMIC-sensitive (eMIC-sens) and eMIC-independent (non-eMIC). For detailed classification rules see Materials and Methods. **D.** Sex comparison for the change in exon inclusion (ΔPSI) between eMIC- and control adult brains. Exon categories as in panel C. **E.** Size distribution of exons regulated by the eMIC domain and all AS exons in *Drosophila* and mouse. Mouse Srrm3/Srrm4 KD data from ^27^. **F.** Proportion of tissue-regulated exons (Supp. Figure 5E) affected by the knockout of the eMIC domain.

To place the eMIC splicing program within the broader AS landscape of *Drosophila*, we analysed published transcriptomic datasets ^28,34,35^ (Table S1) using *vast-tools*, and searched for AS exons with strong tissue-level regulation. Similar to previous reports ^9,10^, we found that neural samples showed the highest prevalence of both tissue-enriched and -depleted exons (Supp. Figure 5E). Interestingly, sensory organs (eye and antenna) displayed a splicing signature that was similar to other neural tissues but with dozens of additional specifically enriched exons, particularly in the eye (Supp. Figure 5E). Remarkably, we found that up to one third of all neural-enriched exons genome-wide are regulated by the eMIC domain (92 eMIC-dep. and 36 eMIC-sens. exons out of 303 neural-enriched exons with sufficient read coverage in our samples; Figure 4F and Supp. Figure 5F), qualifying it as a master regulator of neural-specific splicing in *Drosophila*. Also, AS of the vast majority of these exons were predicted to generate alternative protein isoforms (Supp. Figure 5G), suggesting a prominent role remodelling the neuronal proteome. Finally, we also found numerous muscle-enriched exons, in addition to the previously described splicing singularity of the gonads and sex glands (Supp. Figure 5E and ^9,10^), which were largely not regulated by the eMIC domain (Figure 4F). To facilitate AS research in *Drosophila*, we made these AS profiles publicly available at the *VastDB* website ^36^ (vastdb.crg.eu; example in Supp. Figure 5H).

### eMIC-dependent exons display a unique *cis*-regulatory code

To decipher the regulatory logic of eMIC-dependent splicing, we profiled their sequence features and compared them to other types of exons, including neural non-eMIC-regulated, other AS exons (ASEs), cryptic and constitutive exons (Figure 5A-E, Supp. Figure 6 and Table S2). As shown above, eMIC-dependent exons have a median length of 31nt, strikingly shorter than all other exon types in the *Drosophila* genome (Figure 5A). This short exon size is accompanied by short surrounding introns, closer in size to those neighbouring constitutive exons rather than other types of AS exons (Figure 5A). These very short exon and intron lengths result in low ratio of intron to median exon length (RIME) scores, usually associated with splicing by intron definition ^37^, very similar to the regime of constitutively spliced introns in *Drosophila* and unlike most other alternatively spliced exons ^38^(Supp. Figure 6A). eMIC-dependent exons are further characterised by extremely weak 3′ splice site (ss) regions (Figure 5B), and are also associated with weak 5′ ss but strong upstream 5′ ss compared to constitutive exons, unlike mammalian microexons (Figure 5B and ^20,24^). Additionally, we found a very unique motif architecture in the 3′ ss region: eMIC-exons have long AG exclusion zones, enrichment for UGC motifs close to the 3′ ss, longer polypyrimidine tracts enriched for alternating UCUC motifs, and strong branch-point sequences (BPS; CUAAY motif) (Figure 5C). Intriguingly, the downstream BPS is unusually strong, especially when compared to constitutive exons. Apart from the latter and the weak 5′ ss, all other features are also observed in mammalian eMIC targets (Supp. Figure 6B-F and ^24,26^).

**Figure 5.**
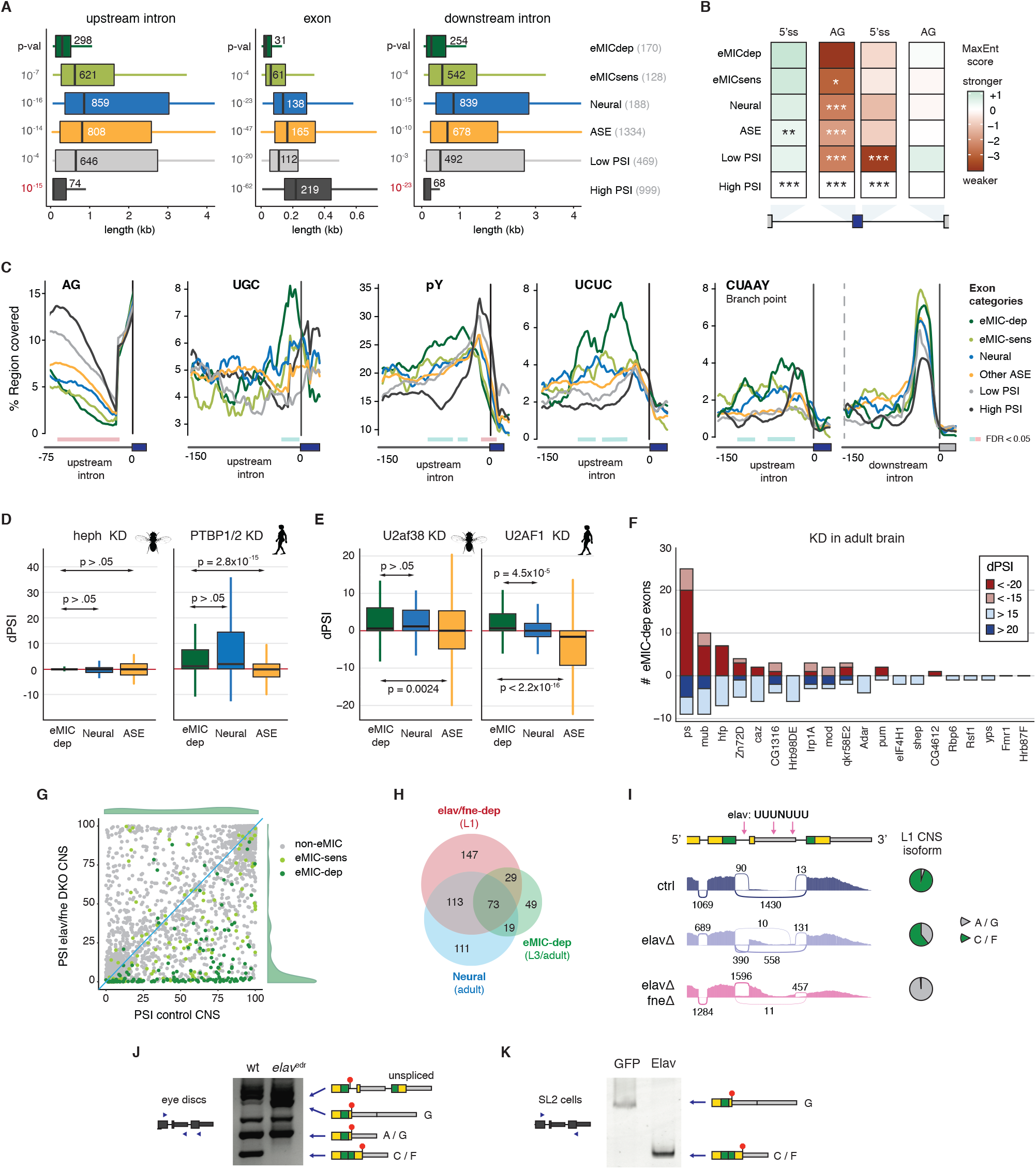
*Cis* and *trans* regulation of eMIC-dependent splicing. **A.** Length of the exon and neighbouring introns for six exon groups, from top to bottom: eMIC-dependent, eMIC-sensitive, Neural, Other AS exons (ASE), cryptic and constitutive exons (Table S2). Box limits represent interquartile ranges; central lines, median values (also indicated with numbers). P-values from Mann-Whitney U tests are shown for the comparison of each class against eMIC-dependent exons. Red font indicates the difference goes in the opposite direction. Number of exons per group is indicated in parentheses. **B.** Maximum entropy scores for the 5′ splice site and AG region, relative to constitutive (High PSI) exons. P-values from Mann-Whitney U tests are shown for the comparison with the eMIC-dependent group: * < 0.01, ** < 0.001, *** < 10^−4^. **C.** RNA maps for motifs enriched or depleted in the intronic regions surrounding eMIC-dependent exons. From left to right: distribution of AG motifs in the 75 nt upstream of the alternative exon start (3′ ss), UGC motif distribution close to the 3′ ss, Polypyrimidine tract profiles for YYYY tetramers (pY) or CU-rich (UCUC/CUCU) tetramers, and profile of the *Drosophila* branch point consensus sequence CUAA[U/C], in both upstream and downstream introns. Length of sliding window: 15 nt for AG and UGC and 27 nt for the others. Regions with a significant difference in the motif coverage (FDR < 0.05) compared to ASE group are marked with a coloured rectangle underneath. **D.** Regulation of eMIC-dependent exons by PTB proteins, quantified as the difference in PSI (ΔPSI) relative to control. Left: *heph* (*Drosophila PTBP1/2/3* ortholog) knockdown in SL2 cells, data from modENCODE. Right: double knockdown of *PTBP1* and *PTBP2* in HEK293 cells, data from ^44^. P-values from Mann-Whitney U tests. **E.** Regulation of eMIC-dependent exons by U2af1. Left: effect of the knockdown of *U2af38* (*Drosophila U2AF1* ortholog) in SL2 cells, data from ^42^. Right: effect of *U2AF1* KD in HEK293 cells, data from ^43^. P-values from Mann-Whitney U tests. **F.** Cross-regulation of eMIC-dependent exons by other RNA binding proteins (RBPs) in fly brains, data from ^45^. In red/blue, number of exons regulated in the same/opposite direction between each RBP and the eMIC domain at two ΔPSI levels. **G.** Effect of the double knockout of *elav* and *fne* on exon inclusion (PSI) genome-wide in the first instar larval (L1) CNS. Data from ^46^. Exons are grouped based on their response to eMIC insufficiency (see Methods). PSI frequency distributions of eMIC-dependent exons in control and mutant CNS are depicted in green. **H.** Overlap between eMIC-dependent exons, neural-enriched exons and Elav/Fne up-regulated exons. The stage of the samples used for the definition of each exon group is indicated. **I.** Sashimi plot of RNA-seq data from L1 CNS upon knockdown of *elav* and *fne*. Pink arrows indicate putative binding motifs of Elav. Data from ^46^. **J,K.** RT-PCR assays of *Srrm234* terminal exons from wild-type (wt) and *elav*-hypomorphic (*elav*^edr^) larval eye imaginal discs (J) and from SL2 cells upon overexpression of Elav (K). Blue triangles mark primer positions. The main splicing products corresponding to annotated isoforms from the *Srrm234* gene (A, C/F, G) are labelled. Non-labelled bands correspond to intermediate splicing products or unspecific amplification that do not differ substantially between samples. wt: wild-type.

We hypothesised that the strong definition of the upstream and downstream splice sites, together with the very short length of the surrounding introns, may facilitate skipping of eMIC-exons outside the neural system, rendering specific repressive *trans*-acting factors unnecessary in *D. melanogaster*. To test this hypothesis we studied the role in *Drosophila* of the main repressor of Srrm4-dependent exons in mammals (Ptbp1)^25,26^. We analysed two RNA-seq datasets upon knockdown of *Hephaestus* (*heph*), the *D. melanogaster Ptbp1/2/3* ortholog, in fly embryos ^39^ and SL2 cells ^40,41^, and found no evidence for a role in repressing the inclusion of eMIC targets, unlike for equivalent experiments in mammalian cells ^25,26^ (Figure 5D and Supp. Figure 7A). To identify potential repressors in an unbiased manner, we also analysed RNA-seq data from knockdown experiments from the modENCODE atlas and others ^40–42^ for dozens of RBPs in SL2 cells (which do not show inclusion of eMIC-dependent exons). This revealed very few factors that might mediate exon repression specifically for eMIC-dependent exons (Supp. Figure 7B,C). Unexpectedly, the top candidate from this analysis was *U2af38* (Supp. Figure 7C), the *D. melanogaster* ortholog of mammalian U2-snRNP auxiliary factor 1, *U2af1*, involved in the recognition of the AG dinucleotide at the 3′ ss, and an interacting partner of the eMIC domain ^24^. This repressive role of *U2af38* was common to other neural exons, but not other ASEs (Figure 5E and Supp. Figure 7C). Remarkably, a similar negative effect on SRRM4-regulated exons was also observed upon knockdown of *U2AF1* in human embryonic kidney HEK293 cells (Figure 5E), and *cis*-regulatory features of eMIC-dependent and U2AF1-repressed exons are notably similar ^43^, suggesting a previously overlooked conserved mechanism across bilaterians.

### Integration of the eMIC splicing program with other regulatory networks

RBPs often cross-regulate each other and co-regulate the splicing of individual AS exons ^41,47–49^. Hence, we searched for *trans*-acting factors whose regulatory programs may overlap with eMIC-dependent splicing by analysing a recent dataset on RBP knockdowns on fly brains ^45^ (Figure 5F and Supp. Figure 7D). Among all factors, the RBP with the largest overlap with eMIC targets was *pasilla* (ps), the mammalian *Nova1/2* ortholog (Figure 5F). This effect is likely due to a direct role in co-regulating eMIC-dependent exons and not indirectly through regulation of eMIC domain expression, since neither *ps* nor any other RBP knockdown from this dataset altered the AS at the 3′ end of *Srrm234* (Supp. Figure 7E).

In contrast, analysis of a recent dataset from first-instar larval (L1) CNS where two members of the ELAV family were knocked out ^46^ gave very different results. *embryonic lethal abnormal vision* (Elav) is an RBP widely used as a neuronal marker that, among other functions, can regulate neuronal AS and APA, and have redundant roles with its paralog *found in neurons* (Fne) ^46,50,51^. Interestingly, the majority of eMIC-dependent exons were completely skipped in L1 CNS upon double depletion of *elav* and *fne* (Figure 5G), and the splicing programs of Elav/Fne and the eMIC domain widely overlapped (p = 8.4 × 10^−142^ hypergeometric test; Figure 5H). In this case, rather than a direct co-regulation of targets, the source of the overlap seemed to be due to the regulation of *Srrm234* last exon selection by the overlapping function of Elav and Fne (Figure 5I-K and Supp. Figure 7F-H). Besides, we identified several putative binding sites for Elav (UUUNUUU motifs) in the 3′ end of *Srrm234* (Figure 5I). Consistently, these two proteins promoted the skipping of the proximal “poison” exon at the 3′ end of *Srrm234* (Figure 5I), but also the selection of the distal polyA site (Supp. Figure 7F). Through RT-PCR assays of the 3′ end of *Srrm234* transcripts in *elav*-hypomorph (*elav*^edr^)^52^ eye imaginal discs, we found that *elav* insufficiency prevented the skipping of the “poison” exon also at this stage and tissue, leading to higher isoform G expression and the complete absence of eMIC expression (Figure 5J). Moreover, heterologous expression in SL2 cells showed that Elav alone is sufficient to promote eMIC expression (Figure 5K and Supp. Figure 7G), and iCLIP (cross-linking and immunoprecipitation) data from fly heads from a recent study ^50^ suggest direct binding of Elav to this region of the *Srrm234* pre-mRNA (Supp. Figure 7H).

In summary, we found that the eMIC splicing program is recruited to neural tissues by the Elav/Fne-mediated regulation of *Srrm234* 3′ end processing and that, within this tissue, it modestly overlaps with splicing networks regulated by other RBPs. Interestingly, we identified several RBPs and transcriptional regulators with alternative isoforms misregulated upon eMIC insufficiency (Table S3), placing the eMIC splicing program within a dense network of neural gene expression regulation, similar to previous results from mammalian model systems ^53^.

### The *Drosophila* eMIC splicing program shapes the repertoire of neuronal ion channels

In line with the Elav-driven expression of the eMIC domain, inclusion of eMIC exon targets increased progressively during embryonic development in neurons but not in glial cells, similar to the pattern observed for *Srrm3/4* targets in mouse (Figure 6A,B and ^26^). Consistent with this observation and with the large overlap with the neural-specific AS program (Figure 4F), the vast majority of eMIC-dependent exons were enriched in all neural tissues: brain, eye, antenna and thoracicoabdominal ganglion (Figure 6C). Still, 15 eMIC-dependent exons were specifically enriched in the eye (Figure 6C), including exons in the ion channels *Otopetrin-like a* (*OtopLa*) and *Chloride channel a* (*ClC-a*), the kinase *retinal degeneration A* (*rdgA*), *crumbs* (*crb*), a key regulator of Notch activity via the Hippo pathway, and its interacting partner *karst* (*kst*). We found that most eMIC-targets were expressed at higher levels in the adult and L1 larval CNS when compared to L3 larval CNS (Supp. Figure 8A-C), likely reflecting the higher proportion of immature neurons in L3 CNS when compared to L1 CNS ^54,55^.

**Figure 6.**
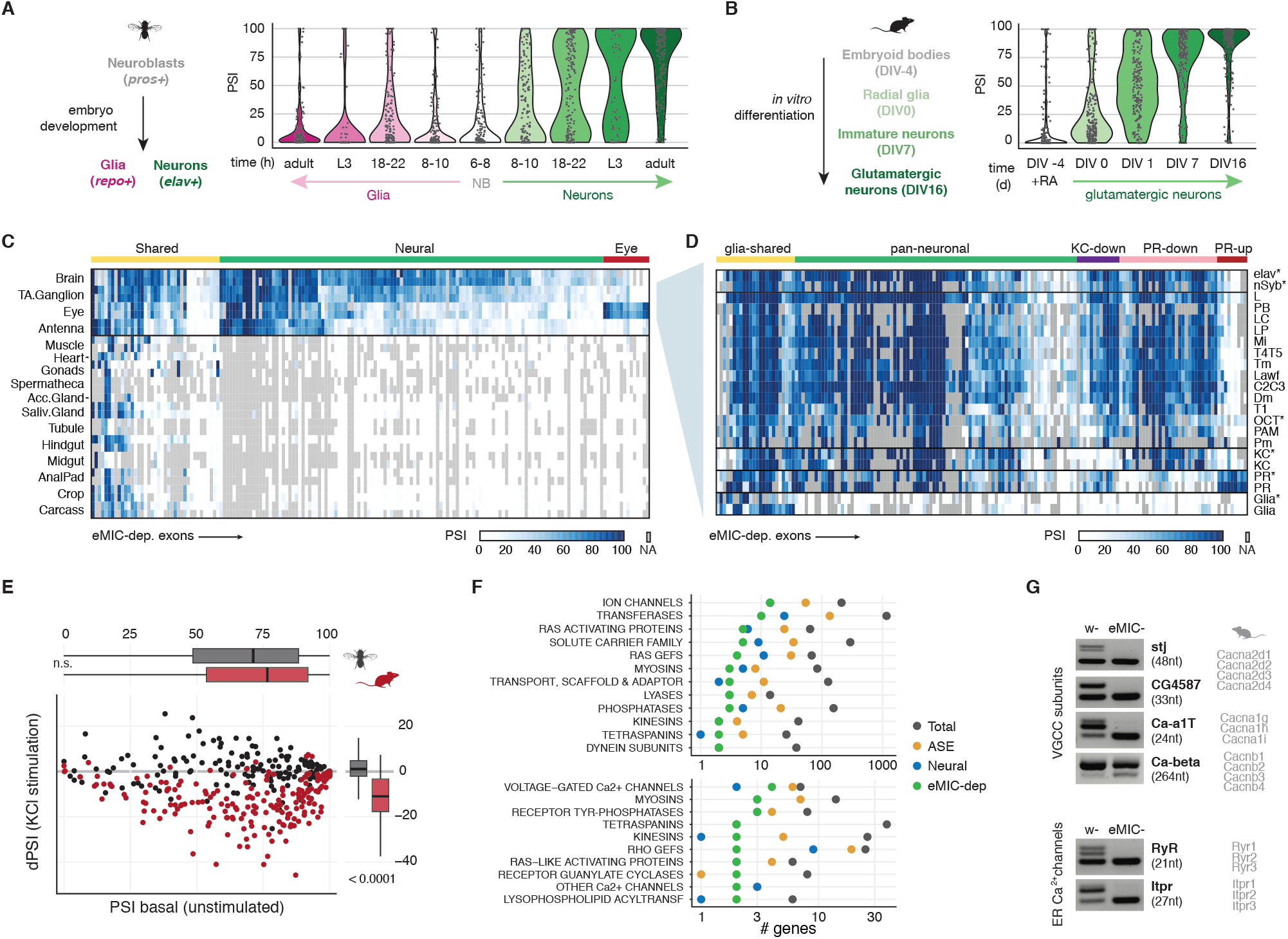
Landscape of the eMIC splicing program across *Drosophila* tissues and cell-types. **A.** Inclusion levels (PSI) of eMIC targets in neuronal and glial cell populations at different times of embryo development as well as larval and adult samples. Data sources in Table S1. NB: neuroblasts, L3: third instar larvae. **B.** Inclusion of mouse eMIC targets along an *in vitro* differentiation time course from embryoid bodies to glutamatergic neurons. Data from ^61^. DIV: days from the onset of differentiation, EB: embryoid bodies, NPC: neural precursor cells. **C.** Heatmap of the inclusion levels of eMIC-dependent exons across adult tissues. TA.Ganglion: thoracicoabdominal ganglion, Acc.Gland: male accessory glands, Saliv.Gland: salivary glands. Data from the FlyAtlas 2 and others (Table S1). **D.** eMIC exon inclusion levels in different neuronal types. Data from ^56^, unless marked with an asterisk (Table S1). KC: Kenyon cell, PR: photoreceptor. Groups are based on the inclusion profile across neural cell types (see Methods for definitions). **E.** Effect of KCl-induced neuronal depolarization on the inclusion of eMIC-dependent exons. *Drosophila* data from ^60^, mouse data from ^22^. P-values from Mann-Whitney U-tests. **F.** Gene groups with more than one member bearing an eMIC-dependent exon. Top: “top-level” gene categories. Bottom: specific gene subgroups. ASE: other alternatively spliced exons. **G.** RT-PCR validations of eMIC-dependent exons in calcium channels from w- and eMIC-heads. On the right, mouse orthologous genes. VGCC: voltage-gated calcium channel, ER: endoplasmic reticulum.

Next, we profiled eMIC-dependent exon expression across cell types in neural tissues, using a recent RNA-seq dataset of cell types in the fly optic lobe ^56^ as well as other datasets of sorted neurons ^57–59^ (Figure 6D, Supp. Figure 8D-H and Table S1). Given that the sequencing depth of some of these samples was too low to robustly quantify exon inclusion, and that eMIC exon inclusion profiles were very similar among closely related cell types (Supp. Figure 8D), we merged related samples to obtain better estimates of exon inclusion (Figure 6D and Supp. Figure 8E). These data revealed several clear patterns. First, we confirmed the neuronal specificity of the splicing program regulated by the eMIC domain, with very few eMIC exons being expressed also in glial cells (Figure 6D and Supp. Figure 8F, “glia-shared” exons). Second, eMIC exons were broadly included across neuronal types but with some degree of variability, with photoreceptors showing the most divergent eMIC exon profile (Figure 6D). As expected, the most photoreceptor-enriched eMIC targets largely overlapped with the set of eye-enriched exons (Supp. Figure 8F), indicating that the eye signature mainly stems from the photoreceptor population. In addition, a fraction of eMIC exons was specifically depleted in photoreceptors and a smaller set was depleted only in Kenyon cells (Figure 6D). These patterns were mirrored by the pan-neuronal expression of the eMIC domain across cell types (Supp. Figure 8H), and by the photoreceptor-specific expression of the newly identified eye-specific *Srrm234* 3′ end isoform (Figure 1C and Supp. Figure 8H), although the variability of the eMIC splicing program followed the same trend as that of all AS exons genome-wide (Supp. Figure 8E,G).

We then looked at the interplay between the eMIC splicing program and transcriptomic signatures of neuronal activity. Mouse Srrm4-regulated exons were shown to decrease inclusion upon sustained KCl-induced neuronal depolarization ^22^. To study if this connection is also present in *Drosophila*, we quantified eMIC exon inclusion using a published dataset of fly brains treated with KCl or activated through optogenetic stimulation ^60^. Unlike the observed effect in mouse, *Drosophila* eMIC exons did not change their inclusion levels upon KCl treatment or optogenetic stimulation (Figure 6E and Supp. Figure 9A). Nonetheless, the behavioural and physiological phenotypes associated with eMIC insufficiency in flies (Figure 3) suggested brain-wide alterations in neuronal activity. Thus, we used these RNA-seq datasets to derive a list of genes up regulated upon KCl or optogenetic stimulation taking into account both end and intermediate timepoints ^60^ (activity-regulated genes, Supp. Figure 9B). We found that this set of genes is overrepresented among the differentially expressed genes in eMIC-brains (7/211, p = 2.4 × 10^−4^, Fisher’s exact test; Supp. Figure 9C and Table S4), further supporting the dysregulation of neuronal activity in these mutants.

To dig into the molecular functions of genes containing eMIC-dependent exons, we next used the gene group classification from FlyBase (Figure 6F). Gene group classification follows a hierarchical organization and thus we focused on both the top and bottom levels, i.e. broad and specific groups, respectively. At the top level, the most numerous category corresponds to ion channels, with 14 genes hosting eMIC exons annotated in this group (Figure 6F, top). Looking at the most specific gene families and complexes, two groups of calcium channels were overrepresented. Firstly, 4 out 7 subunits forming *Drosophila* voltage-gated calcium channels (VGCC) have eMIC-dependent exons: *stj*, *CG4587*, *Ca-α1T* and *Ca-β* (Figure 6F,G). Secondly, 2 out the 3 main intracellular calcium channels are alternatively spliced in an eMIC dependent manner: the ryanodine and inositol-3-phosphate receptors, *RyR* and *Itpr* (Figure 6F,G). These splicing alterations on ion channels, in general, and calcium channels, in particular, may underlie the altered neuronal activity in eMIC-larvae unveiled by our brain imaging experiments.

### Parallel evolution of the fly and mammalian eMIC splicing programs controlling neuronal physiology

The old ancestry of the neural microexon program and the availability of perturbation data from distantly related species make it an appealing case study for the evolution of splicing networks. Hence, we investigated the extent of conservation between the fly and mammalian eMIC-dependent splicing programs. In line with previous studies ^20,24^, ~75% of mouse *Srrm3/4* targets were conserved across tetrapods at the genomic level ^62^ (Figure 7A and Supp. Figure 10A-C). On the other hand, *D. melanogaster* eMIC exons were highly conserved within the *Drosophila* genus, but showed little conservation with other holometabolous insects, such as *Anopheles gambiae* (mosquito), *Tribolium castaneum* (the red flour beetle) and *Apis mellifera* (honeybee) (Figure 7A and Supp. Figure 10D-F). Moreover, fly eMIC exons shared with other holometabolous insects were not always present in closer related species (Supp. Figure 10F), highlighting a higher evolutionary rate of the eMIC splicing program within this clade compared to vertebrates. Nevertheless, we identified high conservation in the flanking intronic sequences of *Drosophila* eMIC targets similar to those of other AS exons in this species (Figure 7B), in line with the proposed regulatory role of these genomic regions. This high conservation was also described for the mammalian eMIC exon program, which showed a particularly high conservation in this region, even when compared to other AS exons (Figure 7B and ^20^).

**Figure 7.**
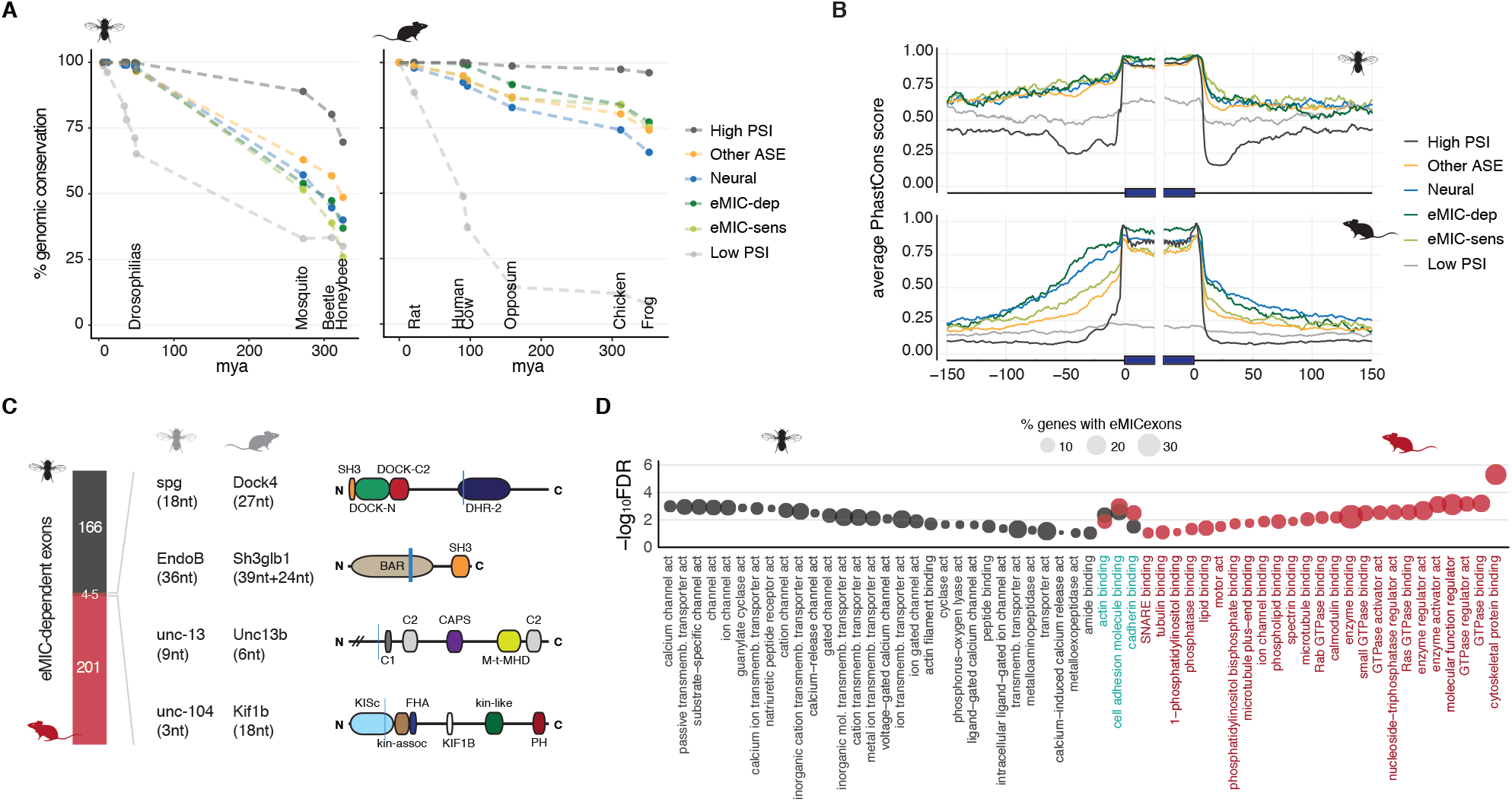
Evolution of the eMIC splicing program in flies and mammals. **A.** Conservation of eMIC-depedent exons at the genome level based on *liftOver*. mya: million years ago. **B.** Sequence conservation of the exonic and flanking intronic regions for six exon groups in mouse and *Drosophila* (Table S2 and Methods). **C.** Overlap between the mouse and fly eMIC splicing programs. For the 4 shared exons, gene name and exon length are indicated in each species. On the right, exon positions are marked within the protein domain scheme with a vertical blue line. **D.** Gene Ontology (GO) terms in the “Function” category that are enriched in *Drosophila* and mouse eMIC targets (black and red, respectively). In blue, terms enriched in both species. FDR: false discovery rate. Circle size represent the percentage of eMIC exon-bearing genes in each GO category.

Remarkably, out of the 157 genes with eMIC exons in *D. melanogaster*, only 19 of them had a mouse ortholog bearing a Srrm3/4-regulated exon, and, out of these, only 4 exons were in the same position as in the fly orthologous gene: *sponge* (*spg*), *Endophilin B* (*EndoB*), *unc-13* and *uncoordinated-104* (*unc-104*) (Figure 7C and Supp. Figure 10G). Moreover, only the exon in *EndoB* could be identified in the genomes of all other studied insect species, favouring a scenario of convergent evolution rather than of common ancestry for the remaining shared eMIC exons. Strikingly, these results thus indicate that the eMIC splicing programs have been nearly completely rewired since their common origin in bilaterian ancestors.

Given this low level of conservation between phyla, we then wondered whether the eMIC domain impacts similar or divergent biological processes in flies and mammals. To avoid biases introduced by gene ontology (GO) annotations in different species, we based our analysis on the more comprehensive human annotation, and performed enrichment analyses of the mouse and fly eMIC targets using GO categories transferred from the human orthologs (see Methods for details). Some GO terms were enriched similarly in mouse and *Drosophila* eMIC targets, indicating a shared bias for genes present in the plasma membrane, cell projections and the synapse, as well as for cytoskeletal proteins (Figure 7D, Supp. Figure 11A). However, most enriched categories were only observed for targets of a single species. The most striking case was the contrasting enrichment for ion channels in the *Drosophila* program and for GTPases in mouse (Figure 7D). Similar results were also obtained using fly GO annotations as reference (Supp. Figure 11B-C) or swapping background gene lists. These results thus suggest that, since the eMIC domain originated in their last common ancestor, each phylum has independently assembled splicing programs that control distinct molecular modules within neurons, which nonetheless ultimately modulate neuronal excitability.

## Discussion

### Control of neuronal physiology by the eMIC splicing program in *Drosophila*

The availability of RNA-seq datasets across fly tissues and developmental stages has uncovered hundreds of splicing decisions that shape their transcriptome and proteome ^9,63^. However, contrary to vertebrate model organisms, most mechanistic studies have focused on the characterization of a handful of individual splice isoforms, lacking a portrait of how the controlled perturbation of broader splicing programs affect physiology. Here, we generated transgenic lines for *Srrm234* isoforms and a loss of function mutant for the eMIC domain, responsible for the regulation of neuronal microexon programs across Bilateria ^24^. We showed that this domain is encoded as an alternative isoform of the pan-eukaryotic *Srrm234* locus that is expressed in neurons due to regulated 3′ end processing by Fne and Elav. These two RBPs have prominent roles on RNA metabolism in neurons and are widely conserved across metazoans ^64–66^, making it good candidates for restricting eMIC expression to neurons also in bilaterian ancestors. In *Drosophila*, we further show that ~28% of Elav/Fne positively regulated exons and one third of all neural-enriched exons depend on the eMIC domain for their inclusion, acting as a master regulator of neuron-specific AS.

Our genome-wide analysis across neural cell-types has highlighted a shared pan-neuronal AS program, with the notable exception of photoreceptors. On a finer level, Kenyon cells in the mushroom body also have a unique splicing signature that is uniform across different studies ^56,58^. Cell-type-specific characterization of the eMIC-regulated splicing program alone mirrored these general AS patterns, placing it as a marker of pan-neuronal identity that nevertheless overlaps with other programs controlling neuron-type-specific AS. Interestingly, a recent study has showed that *Srrm234* (CG7971) expression in mushroom body neurons is required for ethanol-cue-induced memory ^67^. Besides, a different study has identified cycling behaviour of *Srrm234* transcripts in dorsal lateral neurons, potentially connecting this program with the circadian clock ^68^. Here, we show that the splicing alterations in the *Srrm234* eMIC-specific mutant result in an array of neurological-associated phenotypes, most evident of which are locomotion alterations. These, together with the above studies and our brain imaging, sleep and bang-sensitivity results suggest widely pleiotropic effects for this splicing program across the fly nervous system, which could be at least partly explained by the enrichment of eMIC-dependent exons in ion channels and their role in neuronal excitability and function across neuronal populations.

Mis-splicing of ion channels could also underlie the deleterious effect of ectopically expressing the eMIC domain outside the nervous system ^69^, as we describe here for the wing. Additionally, other genes with prominent roles in neuronal function and development are alternatively spliced in eMIC-brains, including genes in key signalling pathways such as Notch (*sno*, *scrib*, *shrb*, *Tsp26A*), BMP (*sax*), Wnt (*spen*), NK-kB (*LRR*), EGFR (*Ptp10D*, *Ptp4E*, *RasGAP1*, *spen*, *Src42A*) and Hippo (*jub*, *crb*) (Table S3). The transgenic constructs generated for *Srrm234* will thus be valuable to dissect the function of this splicing program and the relevance of spatially restricting its expression. It is important to note, however, that our rescue experiments within neurons exposed that quantitative regulation of eMIC expression is particularly sensitive. Hence, more elaborated genetic perturbations might be necessary to fully recover the complex regulation of the *Drosophila Srrm234* locus (Supp. Figure 1) and its function on splicing regulation beyond the eMIC domain, as in the case of the Cwf21 domain in *Srrm234* and the regulation of *Dscam* exon 9 cluster ^70^.

### Evolution of neuronal eMIC-regulated programs in flies and mammals

Comparative transcriptomics across metazoans showed that neuronal microexons originated in bilaterian ancestors driven by the appearance of the eMIC domain, and that neuronal microexons are largely shared within vertebrates ^20,24^. However, conservation rapidly declines outside this group, even in the cephalochordate amphioxus, similar to previous reports on Nova-regulated exons ^71^. Here, by characterizing the full AS landscape regulated by the eMIC domain in *D. melanogaster*, we have confirmed the small overlap between the fly and mammalian programs, with only four exons in equivalent positions within the orthologous genes. Moreover, despite the high conservation within the *Drosophila* genus, eMIC-dependent exons in holometabolous insects show a fast rate of evolution compared to that in vertebrates, with only nine *Drosophila* exons present in all three non-drosophilid insects studied (*A. gambiae*, *T. castaneum* and *A. mellifera*). Notwithstanding, 18 additional eMIC-dependent exons in *Drosophila* and 23 in mouse are present within orthologous genes but at different positions. This recurrence of alternative exons within orthologous genes regulated by the same splicing factor has been previously seen for the *Epithelial Splicing Regulatory Protein* (Esrp) -regulated splicing programs across deuterostomes ^72^, suggesting the presence of hotspots for the evolution of new exons as a common feature in the evolution of splicing programs.

Despite the extensive rewiring of its target exons, the eMIC domain has been associated with neuronal fate expression since its origin in bilaterian ancestors as an alternative isoform of *Srrm234* ^24^. This AS event brought new regulatory capacity to an ancestral regulator similarly to other described cases for transcriptional and splicing regulators ^44,73–78^, ending up in the parallel reassembly of AS programs controlling neuronal function that deploy different modules of the neuronal toolkit in different clades.

## Materials and methods

### Generation of *Srrm234*^eMIC-^ mutant line

The eMIC-null allele for *Srrm234* (*CG7971*) was generated by *GenetiVision* CRISPR gene targeting services. The 650 bp deletion at the C-terminus of the gene was generated using gRNAs *aggtcaaccaaggcggggc* and *gactccggctgttgcgcag* together with donor template harbouring two homology arms flanking a *loxP 3xP3-GFP loxP* cassette (Supp. Figure 2B). Left and right homology arms of the donor were amplified using primers CG7971-LAF3 and CG7971-LAR3, and CG7971-RAF4 and CG7971-RAR4, respectively. Successful deletion and integration of the cassette was validated by PCR and Sanger sequencing, using primers CG7971-outF3 and CG7971-outR4, and LA-cassette-R and Cassette-RA-F, respectively. All primer sequences are included in Table S5. For microscopy experiments, the *3xP3-GFP loxP* cassette was excised by crossing our mutant line with a line expressing Cre-recombinase under a heat-shock inducible promoter.

### Generation of transgenic *Drosophila* lines

Transgenic lines expressing Srrm genes under a 5xUAS promoter were generated at the Francis Crick Institute fly facility. Drosophila Srrm234-A and Srrm234-C isoforms and human SRRM4 open reading frames were sub-cloned from available vectors ^24^. Drosophila Srrm234-I was amplified from fly brain cDNA and verified by Sanger sequencing. This isoform is identical to Srrm234-C but includes an extra intronic sequence that is highly retained in neural tissues. Constructs bear a N-terminal 3xFlag® tag and were cloned into vector pUASTattB and integrated at sites attP40 in Chr2 or attP2 in Chr3, using a line expressing Phi31 integrase from ChrX ^79^. Positive integrants were balanced accordingly to maintain the stock.

### *Drosophila* strains and culture

Flies were maintained at 25°C in 12 h light:12 h dark conditions. A list of the published and unpublished stocks used can be found in Table S5.

### Cloning of *Srrm234* 3′ end minigene and Srrm234 protein isoforms

To generate the *Srrm234* 3′ end minigene, the genomic region encompassing the last 3 exons and corresponding 2 introns (chromosomic region chr3L:1,654,247-1,655,460) was amplified from *D. melanogaster* genomic DNA and cloned in pAc vector. PT1 and PT2 sequences were added upstream to the first exon and downstream to the last exon, respectively ^80^. Additionally, a mutation was performed in the last exon (from nt 148 to nt 164) to introduce a Sp6 motif for detection of the pattern of alternative splicing independently of the endogenous *Srrm234* transcripts. In the case of the tested Srrm proteins, they were cloned in pAc vector with amino-terminal epitopes. pAc vector was a gift from Fatima Gebauer’s lab. Srrm234 proteins isoforms (A, C, I, E) bear a T7-3xFlag^®^ tag, Elav, a 3xFlag^®^ tag, and human SRRM4, a T7 epitope. All primer sequences used for cloning are listed in Table S5.

### Transfection in Schneider S2 cells

400000 Schneider SL2 cells were transfected with 50 ng of plasmid bearing the Srrm234 minigene (when it applies) and 3 ug of Srrm protein expression plasmid (or control) using Lipofectamine 2000 (Invitrogen, following manufacturer’s instructions) and plated in 6-well plates. Cells were collected 48 to 72 hours after transfection. RNA was extracted using RNAspin Mini Kit (GE Healthcare).

### Quantification of gene expression and exon inclusion in fly heads and S2 cells

Flies were snap-frozen in liquid nitrogen and mechanically dissociated by brief vortexing. Heads were separated from the rest of body parts using a mini-sieve system pre-cooled in liquid nitrogen, and stored at – 80 °C. RNA was extracted using NZY Total RNA Isolation kit (MB13402), and cDNA was generated from 250 ng of RNA per sample using SuperScript III (Thermo Fisher Scientific) and anchored oligo-dT primers, following manufacturer’s instructions. RT-PCR assays of alternatively spliced exons were done using primers annealing to the flanking upstream and downstream constitutive exons. Expression levels of *Srrm* constructs were assessed by quantitative PCR using NZYSpeedy One-step RT-qPCR system and detected with a Roche LightCycler® 480 instrument. All primer sequences are provided in Table S5.

### Western blot analysis

Proteins were extracted from SL2 cells pellets using RIPA buffer (20 mM Tris HCl pH 7,5; 150 mM NaCl; 1 mM EDTA; 1% Triton X100; 0,1% SDS; 1 mM DTT and proteinase inhibitors (Roche)). After quantification using BCA (Thermo Scientific), proteins were separated on a 12% SDS-polyacrylamide gel, transferred to a nitrocellulose membrane (GE Healthcare). TBST supplemented with 5% milk was used for blocking and antibodies incubation. The following antibodies were used: Elav (DSHB), HRP anti-rat (Dianova). Membranes were incubated in western blot highlighting Plus ECL solution (PerkinElmer, Inc) before image acquisition using iBRIGHT system. Membranes were staining using Ponceau (Sigma) as a loading control.

### Weight quantification and longevity assay

Flies younger than one day were snap-frozen in liquid nitrogen and weighted in groups of five using a micro-balance with a detection limit of 0.1 mg. Results were provided as average weight by dividing each measurement by 5, a total of 50 flies per genotype were measured. P-values for the difference between genotypes were calculated using Student’s t-tests.

The lifespan at 25°C of 50-100 flies for each sex and genotype was monitored every two days, when flies transferred to a new food vial. Flies that escaped during the transfer were right-censored. P-values for the difference in survival were calculated using the log-rank test.

### Negative geotaxis assay

We separated 3-days-old males and females in groups of 10 flies 24h before the experiment. We transferred the flies to dry 50mL serological pipettes and let then habituate for 2-3 minutes. We video recorded for 1.5 minutes after tapping the cylinders. We manually counted the number of flies crossing each 5mL mark (corresponding to 2.5cm intervals) for each second during 30 s after tapping. We repeated the experiments at least 3 times (30 flies) and calculated mean height positions at each second and nonparametric 95% confidence intervals with 10,000 bootstrap replicates.

### Bang-sensitivity

3-days-old male and female flies were separated in groups of ten individuals 24h before the experiment. The day of the experiment, flies were transferred to clean and dry vials, which were vortexed at maximum speed for 10 seconds. Each vial was the recorded for 2-5 minutes. Recorded videos were manually inspected, number of paralysing flies was quantified and differences between genotypes assessed with Fisher’s exact tests. The time until recovery (i.e. when flies were to upright position) was registered. Flies that did not recover during the duration of the recorded video were right-censored. Statistical differences in recovery time were calculated using Mann Whitney U-tests.

### Sleep and daily activity patterns

Male and female flies were monitored using Trikinetics *Drosophila* Activity Monitors (DAM) using 1-minute bins. Each fly was placed into a glass tube containing 2% agarose and 5% sucrose food. Flies were entrained for 4 days in 12:12 Light:Dark cycles (LD). All the experiments were performed at 25 °C. The sleep and activity parameters of Activity and sleep were analyzed using the MATLAB script SCAMP (https://academics.skidmore.edu/blogs/cvecsey/files/2019/03/Vecsey-Sleep-and-Circadian-Analysis-MATLAB-Program-SCAMP-2019_v2.zip). Activity counts were averaged across 3 days after entrainment for each individual fly. We assayed 16-32 flies per sex and genotype at two ages: recently hatched (3-days-old) and middle age (21-days-old). Statistical differences in maximum sleep duration and total number of activity counts between genotypes were assessed using Mann-Whitney U-tests.

### Larval locomotion analysis

For each experiment, 5 third instar larvae were placed at the centre of a 10 cm Petri dish with 1% agarose and allowed to crawl freely. Dishes were placed on an artist light panel to provide enhanced illumination and recorded from the top at 30 frames per second. Larvae were tracked using BIO (http://joostdefolter.info/ant-research) and output data of larval contours and centres of mass were used for subsequent analysis. Larvae were tracked within a restricted field of view. If a larval contour touched its perimeter (i.e. the larva was exiting this field of view), the larva was not tracked from this timepoint onwards. Where larvae collided, a reliable contour and centre of mass could not be computed – therefore, data during collision was excluded. Larval speed was calculated as the Euclidean distance between the centres of mass of consecutive frames. Mean speed (in pixels per frame) were converted to mm/s by calibrating pixel values to real world values based on the diameter of the behavioural area.

Larval trajectory straightness was determined using the R package ‘trajr’ ^81^. Centres of mass and time values for each larvae were used to create a *trajr* trajectory object. Following this, path straightness was determined using the *TrajStraightness* function, which approximates the efficiency of a directed walk. Straightness indices range between 0 (infinitely tortuous) and 1 (a perfectly straight line). Larval curved body axis was computed using the larval contour output from BIO. As a larva curls into a tighter ball, its contour approximates a circle, which can lead to complications with precise head-tail assignment. Total contour length was calculated as the total Euclidean distances between all contour points. Primary axis circumference was calculated by multiplying the larval primary axis length (defined as being the largest Euclidean distance between pairwise contour points) by pi. A ‘ratio of curvature’ was computed per frame as being the total contour length/primary axis circumference. A ratio of 0.9 and above was taken to indicate larval curling and was used as a threshold to determine the time each larva spent curled.

### Fitness test

We set up F0 crosses using always females of the same genotype and crossing them to males of different genotypes. We quantified the number of flies in the F1 generation based on their sex and the presence of the Sb^−^ marker from the TM3 balancer chromosome. For each genotype and sex, we calculated their relative fitness values as the ratio between their observed allele frequencies and the expected frequencies derived from the Mendelian ratios coming from the mating schemes (0.25:0.25:0.25:0.25). For each cross, we quantified a total number of F1 flies of at least 2,500. Statistical differences in the allele frequencies for each cross were assessed using χ^2^-tests for the average number of flies per replicate.

### Immunostaining

Immunostaining with mouse anti-Fas2 1D4 and anti-brp (nc82) antibodies (Developmental Studies Hybridoma bank) was performed at 1:50 and 1:20 dilutions (respectively) using standard protocols, followed by anti-mouse-Alexafluor-647 (ThermoFisher) or anti-mouse-Cy5 (Jackson Immunoresearch) secondary AB staining, at 1:200 and 1:100, respectively. Nuclei were stained with DAPI and captured with Leica SP5 confocal and processed with FIJI ^82^.

For NMJ staining, third instar larvae were dissected in cold PBS and fixed with 4% paraformaldehyde in PBS for 45 min ^83^. Larvae were then washed in PBS-T (PBS + 0.5% Triton X-100) six times for 30 min and incubated overnight at 4°C with mouse anti-synaptotagmin, 1:200 (3H2 2D7, Developmental Studies Hybridoma Bank, DSHB). After six 30 min washes with PBS-T, secondary antibody anti-mouse conjugated to Alexa-488 and TRITC-conjugated anti-HRP (Jackson ImmunoResearch) were used at a concentration of 1:1000 and incubated at room temperature for 2 h. Anti-HRP is used as a marker to stain neuronal membranes in insects ^84^. Larvae were washed again six times with PBS-T and finally mounted in Vectashield. Images from muscles 6–7 (segment A2-A3) were acquired with a Leica Confocal Microscope SP5. Serial optical sections at 1,024 × 1,024 pixels with 0.4 μM thickness were obtained with the ×40 objective. Bouton number was quantified manually using Imaris 9 software.

### Calcium imaging experiments

Feeding third instar larvae were dissected in physiological saline composed of (in mM): 135 NaCl, 5 KCl, 5 CaCl2-2H2O, 4 MgCl2-6H2O, 5 TES (2-[[1,3-dihydroxy-2-(hydroxymethyl)propan-2-yl]amino] ethanesulfonic acid), 36 sucrose, adjusted to pH 7.15 with NaOH. To prepare the sample for light sheet microscopy, the isolated CNS was embedded in 1% UltraPure^TM^ Low Melting Point Agarose (Thermo Fisher Scientific) in physiological saline at 36 °C and mounted in a glass capillary (1.4 mm inner diameter, 2 mm outer diameter) as previously described ^33^. Once cooled, the agarose was pushed out to expose the embedded CNS outside of the glass capillary. The sample was placed in the imaging chamber filled with physiological saline and spontaneous changes in GCaMP7b intensity were recorded as a proxy for neuronal activity in vGlut-GAL4-expressing neurons in the VNC. Using a Luxendo-Bruker MuVi-SPIM, volumes of either 11 or 22 slices taken every 5-7 μm were imaged with a 4.1 μm light sheet at 0.47 Hz, with an exposure of 20 ms or 2 ms per frame, respectively, and a delay time of 11 ms. Images of 2,048 × 2,048 pixels were taken using a x16 objective.

Only dorsal views were saved and processed manually in FIJI (https://imagej.net/Fiji): images were binned 3 × 3 in XY using BigDataProcessor ^85^, and maximum intensity projections were motion-corrected using StackReg ^86^. Following background subtraction, mean intensities for each half-segment, from the first thoracic to the 8/9th abdominal segment, were obtained using the Multi Measure tool in ROI Manager. Values were imported into MATLAB (R2019b, MathWorks) and ΔF/F = (F(t) - F_0_) / F_0_ traces were calculated, where F(t) is the fluorescence intensity at a given time point and F_0_ is the mean fluorescence intensity in a manually defined window lacking any spontaneous activity, spanning 10 frames (~ 21.3 s).

To identify activity bouts, ΔF/F data was averaged across segments, smoothened using a low-pass filter, and local maxima were identified using the “findpeaks” function in MATLAB. Smoothened traces were obtained using a Short-time Fourier Transform (FT), followed by an inverse FT with a low-pass filter. In brief, data was split into fragments of 100 frames (~ 3.5 min) overlapping by ½, each fragment was windowed using a Hamming function, its discrete FT computed, and the resulting magnitudes and phases were used to reconstitute the data from the lowest 5 frequency components.

To identify the number of waves and their mean amplitudes, ΔF/F data was averaged across segments and “findpeaks” was used to extract the locations and amplitudes of local maxima in each sample. Peak amplitudes were measured from baseline. Next, ΔF/F data from 3 frames flanking each peak location (resulting in 7 frames = 14.9s) was extracted from each segment, and the location of the maximum peak identified, together with its width at half-height. For each wave, the slope of a linear fit for peak locations as a function of VNC segment was used to determine its type, with a negative slope identifying a forward wave and a positive slope identifying a backward wave. Peak widths were used to determine the starting and ending points of a wave.

One-sided activity was assessed by measuring absolute differences in intensity between each left-right segment pair. Values represent the percentage of recording time in which the absolute difference is larger than a predefined threshold of 0.7 ΔF/F on average per sample. The threshold was established by comparing visually observed one-sided activity to noise in two example samples. Matlab scripts are available at https://github.com/snznpp/eMIC_GCaMP.

### Single-cell RNA-sequencing

50 full central nervous systems (brain lobes and ventral nerve cord) were dissected from L1 larvae 1-2 h after hatching for control w^1118^ and eMIC-larvae in ice cold Schneider’s medium and transferred into an embryo dish with 1x DPBS on ice. Total dissection time did not exceed 2 hours. After dissection, samples were transferred into 0.5 mL vials containing dissociation solution (DPBS with papain 0.2 U/mL, collagenase 0.1 mg/ml and BSA 0.04%). Brains were briefly centrifuged at 4 °C, resuspended in 80 μL of fresh dissociation solution and left dissociating for 30-40 minutes at 25 °C shaking at 1000 rpm. Every 5 minutes, tissue samples were triturated using a 200 μL pipette tip (50-80 times). After dissociation, cells were passed through a 20 μm Flowmi Cell strainer and centrifuged at 2000 rpm for 5 minutes at 4 °C. Dissociation solution was removed and substituted by DPBS + 0.04% BSA. 10 μL of dissociated cells were used to determine cell yield by using Hoechst staining under a fluorescent stereomicroscope and a C-Chip Neubauer. Samples were adjusted at a concentration of 1200 cells/μL for loading into Chromium single-cell chip. scRNA-seq libraries were prepared using the Chromium Single Cell 3’ Library and Gel Bead Kit v3 according to manufacturer’s instructions. Sequenced libraries were processed using Cell Ranger v2.0.0 provided by 10x Genomics, resulting in 29,554 cells with 1232 unique molecular identifiers (UMIs) and a median of 693 genes per cell for control sample, and 29942 cells with 1205 UMIs and a median of 679 genes per cell for the eMIC-mutant sample.

### Single-cell RNA-seq analysis

Sequencing data was analysed using Seurat 4.0.0 R package. Cells with less than 5% mitochondrial genes and a unique feature count between 50 and 2500 were selected for downstream analysis, leaving 19400 cells in the control and 18268 cells in the mutant sample. Data was normalised with a log scale factor of 10000 and 2000 highly variable features were selected according to the R package’s developer recommendation. After inspection of Elbow plots, Jackstraw plots and principal components heatmaps, 20 principal components were selected to explain the variability of the data. To subdivide the datasets into clusters, the Seurat command FindNeighbours with a HD.Dim 20 and FindClusters with a resolution of 0.5 were used. Finally, the dataset with reduced dimensionality was visualised using a UMAP plot, which led to the generation of 16 clusters in both the control and the mutant datasets. Data integration for control and mutant samples was performed by identifying cell types based on marker and common anchor genes as implemented by the IntegrateData function of Seurat, producing a single integrated Seurat Object. Within the integrated data, common cell types were identified based on known marker genes from visualizations using the FeaturePlots Seurat command.

### Bulk RNA-sequencing

Wild type (OreR) and Srrm234^eMIC−^ mutant flies were raised at 25°C and 12h:12h dark:light cycle. Late female and male L3 instar larvae and 24 h female and male adults were collected separately and their brains (> 20 per sample) dissected in PBS 1X and stored in RNALater (Qiagen, Venlo, Netherlands). Total RNA was isolated using RNeasy Mini Kit (Qiagen) and RNA quality was checked using Bioanalyzer (Agilent). A total of eight strand-specific Illumina libraries were prepared and sequenced at the CRG Genomics Unit. An average of 80 million 125-nt paired-end reads were generated for each sample.

### Visualization of genomics data

bigWig files with 3′ seq data of elav and fne mutants and bedGraph files with elav iCLIP data were downloaded from GEO (accession numbers GSE155534 and GSE146986, respectively). 3′ seq and CAGE data from different tissues were also downloaded from GEO dataset GSE101603 and modENCODE ^9^. Data was visualized using IGV (Integrative Genomics Viewer) web browser ^87^. For visualization of splice junctions (sashimi plots) we mapped reads using STAR ^88^ and generated sashimi plots for specific genomic regions using *ggsashimi* ^89^.

### Alternative splicing analysis of eMIC KO brains

AS analysis was performed using PSI (Percent Spliced In) values calculated with *vast-tools* v2.5.1 ^36^ for dm6 (VASTDB library: vastdb.dme.23.06.20.tar.gz), filtering out events with very low number of mapped reads (minimum quality score of LOW or higher). Global analysis of AS changes in eMIC mutant brains was done using a change in average PSI (ΔPSI) of 20 between mutant and control brains and requiring a minimum ΔPSI of 15 between genotypes for each sex independently, in either adult brains or third instar larval central neural systems. Only AS events mapping to the coding sequence were considered.

For exon classification based on the regulation by the eMIC domain we used the following criteria:

- ‘eMIC-dependent’: cassette exons that have substantially lower inclusion in eMIC-mutant brains. We used the previous criteria ΔPSI ≤ −20 in either adult or larval samples and minimum ΔPSI ≤ −15 for each sex. We also added exons with a ΔPSI ≤ −10 in both sexes if the inclusion level in the knockout (KO) samples was very low (PSI ≤ 1).
- ‘eMIC-sensititve’: exons not included in the above group that show a ΔPSI ≤ −10 in both sex-paired comparisons between wild-type and KO samples, either for adult or larval samples. We also considered ‘eMIC-sensitive’ exons those enhanced by the overexpression of human SRRM4 in SL2 cells ^24^ with a ΔPSI ≥ 15 and that were not considered ‘eMIC-dependent’.
- All other exons were considered eMIC-independent as long as they had sufficient coverage in our brain samples (minimum *vast-tools* score of LOW).

eMIC-dependent exons that were differentially spliced between males and females were defined as those with |ΔPSI| ≥ 20 between wild-type and eMIC-knockout brains in either sex, that have a |ΔPSI| ≥ 20 between male and female samples in control or mutant brains.

### Quantification of Srrm234 last exon usage

We quantified alternative last exon usage at the *Srrm234* locus as the proportion of reads mapping to each of the three possible exon-exon junctions from the same common donor in the first eMIC-encoding exon to each possible acceptor site. The number of mapped reads for each of the three junctions was obtained from the eej2 output of *vast-tools*. The shared donor corresponded to donor 20, and the three quantified acceptors were 21, 22 and 23 from locus FBgn0035253.

### Genome-wide cell and tissue type specific splicing patterns

We collected publicly available RNA-seq data from fly adult tissues from different sources (Table S1), quantified PSI values using *vast-tools* v2.5.1 ^36^, and grouped exons based on their inclusion profiles. For this, we grouped tissues into eight groups: neural, sensory, muscle, digestive tract, salivary glands, ovary, testis and sex glands. Next, to define tissue-enriched exons we required a ΔPSI ≥ 25 between that tissue and the average of all other tissues and ΔPSI ≥ 15 between that tissue and the maximum PSI across all other tissues. Similar analyses but changing the direction of ΔPSI were done to define tissue-depleted exons. For the case of neural and sensory tissues, we excluded each other from the “all other tissues” group, given their partially overlapping cell-type composition. AS analysis across neural cell types was performed using a recently published dataset containing over 100 samples ^56^ together with other independent samples (see Table S1 for full list of samples). We assessed global patterns of AS exons by looking at exons with a PSI range ≥ 20 across neuronal and glial samples.

### Tissue- and cell-type-specific profiles of eMIC-dependent exons

We classified eMIC-dependent exons into three groups based on their inclusion profiles across tissues: (i) ‘Shared’: exons that were not enriched in neural tissues with a PSI ≥ 40 in any non-neural tissue or with a ΔPSI ≤ 15 when comparing neural and non-neural samples, (ii) ‘Eye’: eye-enriched exons with a ΔPSI ≥ 20 between eye and other neural tissues (brain and thoracicoabdominal ganglion), and (iii) ‘Neural’: all other eMIC-dependent exons showing neural enrichment, as described in the previous section.

Analogously, we classified eMIC-dependent exons into five groups based on their inclusion profiles across neural cell types: (i) ‘Glia-shared’: exons with a PSI ≥ 50 in any glial sample, (ii) ‘PR-up’: exons with a ΔPSI ≥ 25 between photoreceptors and other neuronal types, (iii) ‘KC-down’: exons with a ΔPSI ≤ −25 between Kenyon cells and other neuronal types, (iv) ‘PR-down’: exons with a ΔPSI ≤ −25 between photoreceptors and other neuronal types, and (v) ‘pan-neuronal’: the rest of eMIC-dependent exons with neuronal enrichment. For calculating sample-to-sample distances based on the inclusion profile of eMIC-dependent exons or all AS exons across neural types, we used (1 – pearson correlation) as the clustering distance between samples.

### AS analysis of mammalian exons

We quantified AS genome-wide using the same method as described above for *D. melanogaster*, i.e. *vast-tools* v2.5.1 for mouse mm10 (VASTDB library: vastdb.mm2.23.06.20.tar.gz) and human hg38 (VASTDB library: vastdb.hs2.23.06.20.tar.gz) genome assemblies. Definition of mouse eMIC-dependent and eMIC-sensitive exons was based on publicly available data of the double knockdown (KD) of *Srrm3* and *Srrm4* in N2A cells ^27^ and the knockout (KO) of *Srrm4* in mouse hippocampus and cerebral cortex ^21^. We classified exons as ‘eMIC-dependent’ if they had an average ΔPSI ≤ −20 between mutant and control samples and minimum PSI range ≤ −15 between conditions; or had an average ΔPSI ≤ −10 between mutant and control samples and maximum PSI ≤ 1 in mutant samples. ‘eMIC-sensitive’ exons were those that were not included in the previous group and that had a ΔPSI ≤ −15 and PSI range ≤ −5 between *Srrm3/4* KD and control samples in N2A cells; or that had a ΔPSI ≤ −10 between *Srrm4* KO and control samples in both hippocampus and cortex. Exon groups used as controls: Neural, ASE, LowPSI and HighPSI were defined as described in ^24^ (and see below) and are included in Supp. Table S2. Definition of human eMIC-dependent exons was based on overexpression data of SRRM4 in HEK293 cells ^24,90^: we required a ΔPSI ≥ 40 between overexpression and control and |ΔPSI| ≤ 10 between replicates. Neural and ASE control groups were also taken from ^24^ and included in Table S2.

### Analysis of exon/intron features

Analysis of the *cis*-regulatory code associated with eMIC-dependent splicing was done by comparing the eMIC-dependent and eMIC-sensitive exon sets to four control exon sets (Table S2). We defined these groups based on their inclusion profiles across the eight tissue groups defined above: (i) ‘HighPSI’: highly included exons with a minimum PSI > 90 across all tissues with sufficient read coverage (*vast-tools* score LOW or higher), (ii) ‘LowPSI’: lowly included exons with a maximum PSI < 10 across all tissues with sufficient read coverage, (iii) ‘Neural’: Neural-enriched exons (as defined above) not regulated by the eMIC domain, and (iv) ‘ASE’: alternatively spliced exons with sufficient read coverage in at least in 3 different tissues and that are alternative (10 ≤ PSI ≤ 90) in at least 25% of all samples. HighPSI exons were down-sampled to 1,000 exons by random selection.

We calculated the strength of the donor and acceptor splice sites (5′ss and 3′ss/AG, respectively) according to maximum entropy score models. To test for differences in the median of the scores for each feature we used Mann-Whitney U tests. To generate the RNA maps in Figure 5 and Supp. Figure 6 we used *rna_maps* function from *Matt* v1.3 ^91^ using sliding windows of 27 nucleotides. Polypyrimidine tract were searched as YYYY tetramers, CU-rich motifs included the motifs UCUC and CUCU, and the *D. melanogaster* branch point consensus sequence used was CUAAY. For estimating FDR values in the RNA maps we used a permutation test using 1,000 permutations and a threshold of FDR < 0.05 as implemented in *Matt*. Ratio of Intron to Mean Exon length (RIME) was calculated for the skipping isoform of each exon. The total length of the intron was calculated as the sum of the alternative exon and the upstream and downstream introns, and mean exon length was calculated for the adjacent constitutive exons.

### Protein impact prediction

Exons detected by *vast-tools* that mapped to coding sequences were classified following the description in ^20^, as provided in *VastDB* (version 2.2 of protein predictions). In brief, exons were predicted to disrupt the coding sequence (CDS) if their inclusion or skipping would induce a frameshift in their open reading frame (ORF) or if they would induce a premature stop codon (PTC) predicted to be targeted by nonsense-mediated decay (NMD) or to truncate the protein by more than 300 amino acids or more than 20% of the reference isoform. The rest of CDS-mapping exons are predicted to preserve the transcript coding potential.

### AS analysis of RNA-binding protein perturbations

Published data of CNSs from *elav* and *fne* mutant first instar larvae was analysed using *vast-tools* v2.5.1. We considered ‘elav/fne-dependent’ exons those with an average ΔPSI ≤ −20 and a PSI range ≤ −15 between elav/fne double-KO and control samples. We analysed the effect of knocking-down a collection of RNA binding proteins (RBPs) in Drosophila SL2 cells using data from modENCODE and REF-U2af. We processed this data with *vast-tools* and quantified the effect of each KD on different types of AS exons: eMIC-dependent, eMIC-sensitive, Neural and other AS exons -ASEs (defined above). We tested the difference in the ΔPSI of each knockdown on eMIC-dependent exons and compared it to the effect on ASEs. We compared the differences between the two exon groups for each RBP using Mann-Whitney U-tests correcting for multiple testing with the Benjamini-Hochberg method.

To test the overlap with other splicing programs controlled by different RBPs we used a publicly available dataset on RBP KDs in the fly brain (^45^, Table S1). We calculated the ΔPSI between each RBP KD and the control GFP-KD sample for all eMIC-dependent exons and considered as regulated those with an |ΔPSI| ≥ 20 or 25. For this and the SL2 dataset we assessed the overall KD efficiency of each RBP by calculating the fold-change in gene expression for each sample (Supp. Figure 7B,D)

The same approach was followed to calculate the effect of knocking down the human U2af38 and heph orthologs in HEK293 cells: U2AF1 and PTBP1/2, respectively, and for mouse Ptbp1/2 KD in embryonic stem cells. All data sources are included in Table S1 and necessary files to run *vast-tools* are indicated in ‘code availability’ section.

### Differential gene expression analysis

Gene expression quantification was done using *vast-tools* v2.5.1. For analyses of eMIC mutant brains, we considered female and male samples as replicates given their high degree of similarity. To quantify differential gene expression (DGE), raw gene counts calculated with *vast-tools* for wild-type and mutant adult brains were used as input for edgeR ^92^. DGE was estimated using the likelihood ratio test (LRT) between genotypes taking the sex factor into consideration (~ genotype + sex). False Discovery Rates were calculated using the Benjamini-Hochberg method. DEGs with a FDR ≤ 0.1 and fold-change ≥ 1.5 in adult mutant brains are listed in Table S4. Statistical assessment of the number of activity-regulated genes (ARGs) that are differentially expressed in eMIC mutant brains was determined using Fisher’s exact test.

### Analysis of neuronal-activity DGE and AS

We assessed the effect of sustained neuronal activation on gene expression and AS using a publicly available dataset using two stimulation paradigms: KCl-induced depolarization and optogenic activation ^60^. We assessed DGE using the ‘exact test’ method as implemented by edgeR comparing each time point with its corresponding un-stimulated sample for each of the two stimulation paradigms. FDR were calculated using the Benjamini-Hochberg method. We considered activity-regulated genes (ARGs) those with an FDR value lower than 5% and with a fold-change ≥ 1.5 compared with control samples at any time point. PSI values for all exons mapping to coding sequences genome-wide were calculated as described above.

### Gene Group classification of genes with AS exons

We classified genes harbouring AS exons into functional groups according to the classification directly downloaded from FlyBase gene groups (“gene_group_data_fb_2020_01.tsv”). This classification is hierarchical so we looked at two levels: the “bottom” categories with very specific gene subgroups and the “top” level group for each subgroup. We did this analysis for three types of AS exons: eMIC-dependent, neural non-eMIC-regulated and other AS exons. We plotted both the number of genes per group and their proportion of the total number of group members.

### Sequence conservation of the genomic region surrounding AS exons

phastCons data files were downloaded from UCSC Genome Browser server. For mouse, we used phastCons data from an alignment of 60 vertebrate genomes and for *D. melanogaster* we used data from the alignment of 27 insect species. We averaged phastCons scores with respect to the start and end of our six previously defined exon groups for both species. We removed overlapping gene elements that may distort the signal such us upstream and downstream exonic sequences if these are closer than 150 bp, and the first 10 nt of the upstream intron and the last 30 nt of the downstream intron to avoid signals coming from the splicing donor and acceptor sites.

### Exon-level conservation with liftOver

To look at genomic conservation of AS exons we used UCSC liftOver. For *D.melanogaster*, we used liftOver chain files for dm6 genome annotation with all available species. For mouse, we used representative species covering a similar time window as the available species for Drosophila, i.e. rat, human, cow, opossum, chicken and frog. We defined 4 levels of exon conservation: i. the entire region is missing from the chain file in the second species (including adjacent constitutive exons), ii. adjacent exons can be liftOvered but not the AS exon region, iii. the genomic region encompassing the AS exon can be liftOvered but none of the exon splice sites are present, and iv. at least one of the splice sites from the AS exon is present (either the acceptor or the donor sites). We considered exons to be conserved at the genome level if they belong to the last group.

### Overlap of eMIC-dependent splicing programs between fly and mouse

We defined orthology relationships between mouse and fly using DIOPT (http://www.flyrnai.org/diopt). For conservation analysis of eMIC-dependent exons we first assigned gene orthology requiring a DIOPT score ≥ 2 and “best score” in at least one direction. Orthologous genes with eMIC-dependent exons in both species were then pairwise aligned using MUSCLE ^93^. Exons were considered orthologous if they are in the same intron in both species, i.e. having the same the same adjacent exons.

### Gene Ontology analysis

To avoid biases derived from species-specific gene annotations, we followed a strategy based on orthology relationships to assess Gene Ontology (GO) enrichment. First, we generated two lists of genes for each species: eMIC-dependent exon containing genes and a background gene set. The background list contained those genes with a minimum expression level similar to eMIC-containing genes in the datasets used for calling of eMIC-dependency: cRPKM ≥ 5 in adult fly brains or cRPKM ≥ 3 in N2A cells. These gene lists were then transformed to their human orthologs with similar criteria as specified above: DIOPT score ≥ 2 and “best score” required (“best-reverse” score not sufficient). We then run GO enrichment analysis using GOrilla ^94^ with the ‘two unranked lists of genes’ module using the human annotation. We joined the output obtained using the human orthologs of mouse and fly genes for each category (Process, Function and Component) and visualized them using ggplot2. Similar results were obtained when swapping the human ortholog background lists generated using either the fly or mouse data. Finally we also performed the same analysis using the fly orthologs of mouse genes and running GOrilla using the *D. melanogaster* GO annotation to explore GO terms associated with fly-specific biological processes. All GO terms enriched for the Function and Component categories are represented in Figure 7D and Supp. Figure S10, and all enriched GO terms are listed in Table S3.

### Code availability

All software used to analyse the data is publicly available and indicated in the Methods section. VASTDB files to run *vast-tools* are available to download for each species (https://github.com/vastgroup/vast-tools) as indicated in the Methods section. Custom code to generate figure plots is available upon request.

### Data availability

RNA-sequencing data was submitted to SRA and included within the “Origin and Evolution of Neural Microexons” BioProject (PRJNA474911). All other RNA-Seq samples used in this study are publicly available and listed in Table S1. Quantifications of AS and gene expression across most samples included in this study can be found in the repository vastdb.crg.eu.

## Acknowledgements

The research has been funded primarily by the European Research Council (ERC) under the European Union’s Horizon 2020 research and innovation program (ERC-StG-LS2-637591 to MI), the Spanish Ministerio de Ciencia (BFU2017-89201-P to MI), the Francis Crick Institute, which receives its core funding from Cancer Research UK (FC001594), the UK Medical Research Council (FC001594), and the Wellcome Trust (FC001594), a European Research Council (ERC StG “EvoNeuroCircuit” 802531) to LPG, and the ‘Centro de Excelencia Severo Ochoa 2013-2017’(SEV-2012-0208). ATM held an FPI-Severo Ochoa fellowship and a travel grant from Boehringer Ingelheim Fonds. RJVR held a Boehringer Ingelheim Fonds PhD fellowship. Work in SK lab was funded by the NIH R01 grant (R01GM122406) and work in MS lab by BBSRC. We acknowledge the support of the CERCA Programme/Generalitat de Catalunya and of the Spanish Ministry of Economy, Industry and Competitiveness (MEIC) to the EMBL partnership. For the purpose of Open Access, the author has applied a CC BY public copyright licence to any Author Accepted Manuscript version arising from this submission. We would like to thank Alessando Ciccarelli at the Light Microscopy STP, Amy Strange, Luke Nightingale and Joost de Folter at the Scientific computing STP and the Fly Facility at The Francis Crick Institute for their support and assistance in this work, and Chris Rookyard for his help and advice on calcium imaging analysis.

## Author contributions

ATM performed all bioinformatics analyses and experiments not specified below and generated fly stocks. SP performed and analysed calcium-imaging experiments. SB performed experiments with S2 cells. IA performed immunostainings and molecular biology work. AAv performed and analysed scRNA-seq experiments. RJVR analysed larval locomotion behaviour. CP did NMJ experiments under supervision of JYR. AAl, VM and IUH performed molecular biology work under supervision of FC, JYR and MS. AMA and SK supervised daily activity, longevity and larval locomotion assays. MI and LPG supervised the study and provided resources. ATM, LPG and MI wrote the manuscript with input from all authors.

## Supplementary Material

**Table S1.** List of RNA-seq samples from *Drosophila*, human and mouse, analysed in this study.

**Table S2.** List of eMIC-dependent exons in *Drosophila* and mouse, and classification of the fly exons according to their profile across neural cell types.

**Table S3.** Gene Ontology (GO) terms enriched in *Drosophila* and mouse eMIC-dependent exons using the human GO annotation; and signalling- or gene regulation-related genes with eMIC-dependent exons in *Drosophila*.

**Table S4.** Differentially expressed genes in *Drosophila* eMIC-adult brain.

**Table S5.** List of primers and fly stocks used in this study.

**Video S1.** Self-righting defects in eMIC-male adult flies.

**Video S2.** Body posture and locomotion defects in *elav>Srrm234-I*;eMIC-male adult flies.

**Video S3.** Wave amplitude of control and eMIC-larval central nervous system during fictive locomotion experiments.

**Supplementary Figures 1–11.**

---

**Supplementary Figure 1.**
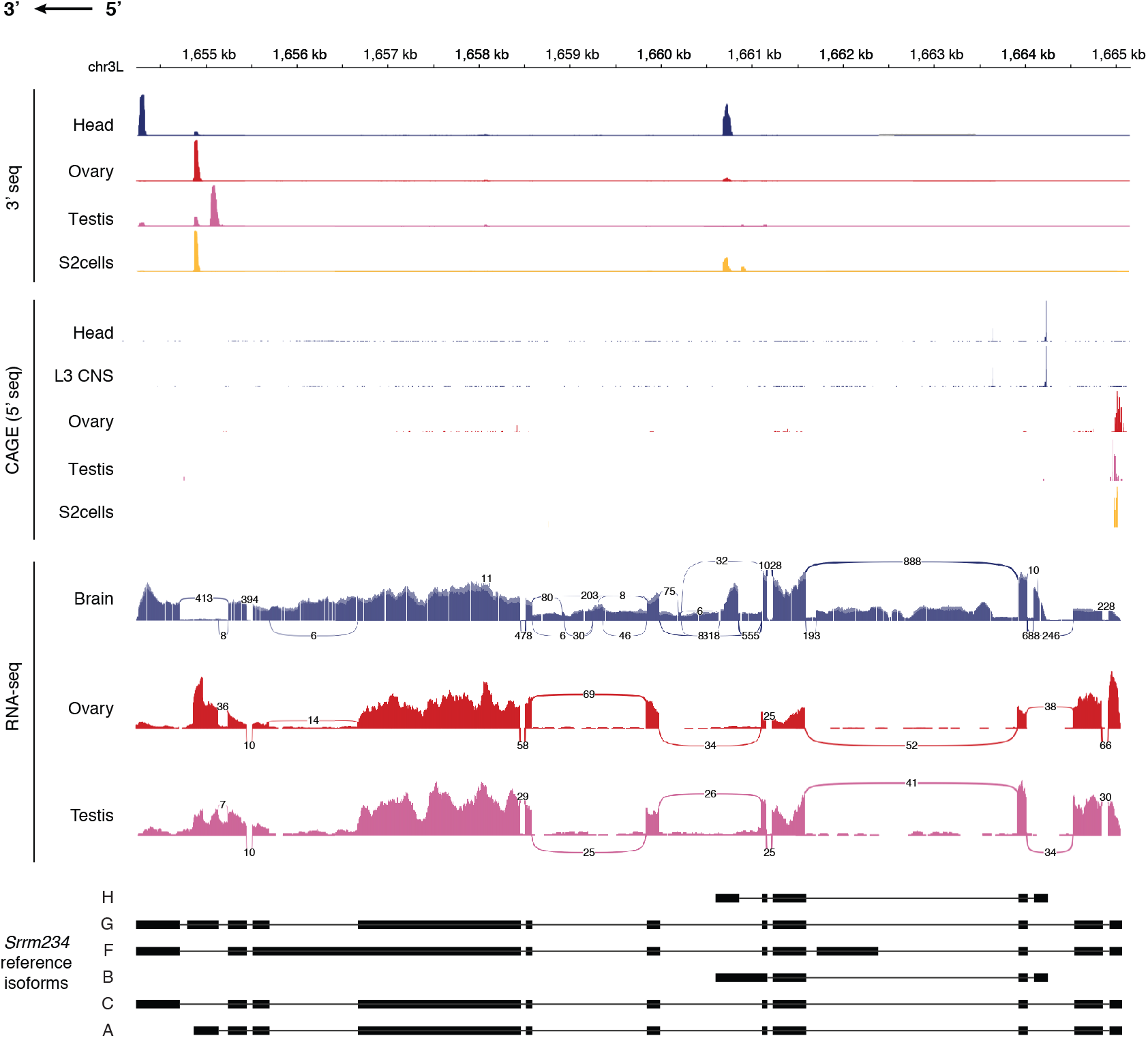
*Srrm234* alternative isoforms across tissues. **A.** Transcriptomic data mapping at the Srrm234 gene. Top, 3′ seq data from ^95^. Middle, CAGE-seq (cap analysis gene expression) data from ^9^. Bottom, RNA-seq data from Fly Atlas 2 ^28^.

**Supplementary Figure 2.**
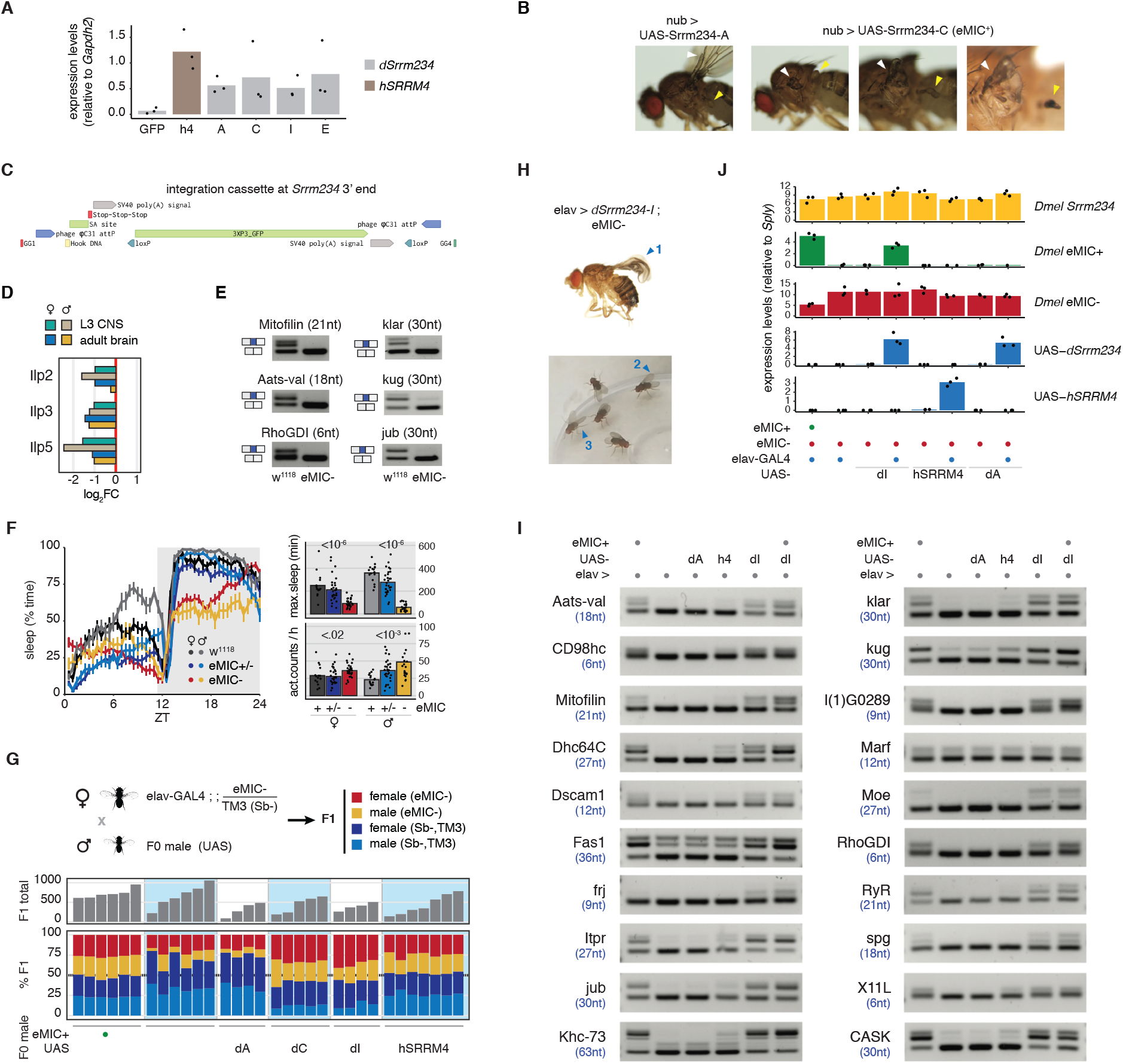
Phenotypic characterization of flies lacking or overexpressing the eMIC domain. **A.** Expression levels as quantified by qPCR of heterologous expressed *Srrm234* constructs (*Drosophila* isoforms A, C, I, E and *hSRRM4*) in SL2 cells, relative to the housekeeping *Gapdh2* gene. **B.** Representative wings (white arrows) and halteres (yellow arrows) of flies expressing *Srrm234*-derived transgenes under the control of nub-GAL4 (*nubbin*) driver lines. **C.** Integration cassette introduced at the 3′ end of *Srrm234* replacing the endogenous sequence that encodes for the eMIC domain (region delimited by guide RNAs in Figure 2A). **D.** Expression levels of are neuronally secreted insulin-like peptides (Ilps) in *Drosophila*, in control and eMIC-L3 CNS and adult brain. **E.** RT-PCRs of alternatively spliced exons in fly heads for control (w^1118^) and eMIC-flies. In brackets, exon lengths. **F.** Sleep patterns of 21 day-old flies in 12h light – 12h dark cycles. Left, average time flies spend sleeping (inactive for >= 5min) at different times of the light:dark cycle. ZT: Zeitbeger Time (switch from light to dark conditions), vertical lines: standard error of the mean. Top right, maximum sleep episode during the night. Bottom right, total number of activity counts per hour during the night. P-values from Mann-Whitney U tests comparing with w^1118^ controls. **G.** Top, mating scheme for the determination of relative fitness values presented in Figure 4B. Sb: *stubbl*e. Middle, total number of flies quantified per experiment. Bottom, allele frequencies in the F1 generation. **H.** Deleterious effects of overexpressing dSrrm234-I pan-neuronally in the eMIC-background: failed wing expansion (1) and open positioning of wings (2) and legs (3). **I.** RT-PCRs from female fly heads of candidate eMIC-sensitive exons. dI: *Drosophila* Srrm234-I, h4: human SRRM4, dA: *Drosophila* Srrm234-A. **J.** Expression levels as quantified by qPCR of *Srrm234* gene, each of the two alleles (eMIC+ and eMIC-), and UAS-transgenes in female fly heads, relative to the housekeeping *Sply* gene.

**Supplementary Figure 3.**
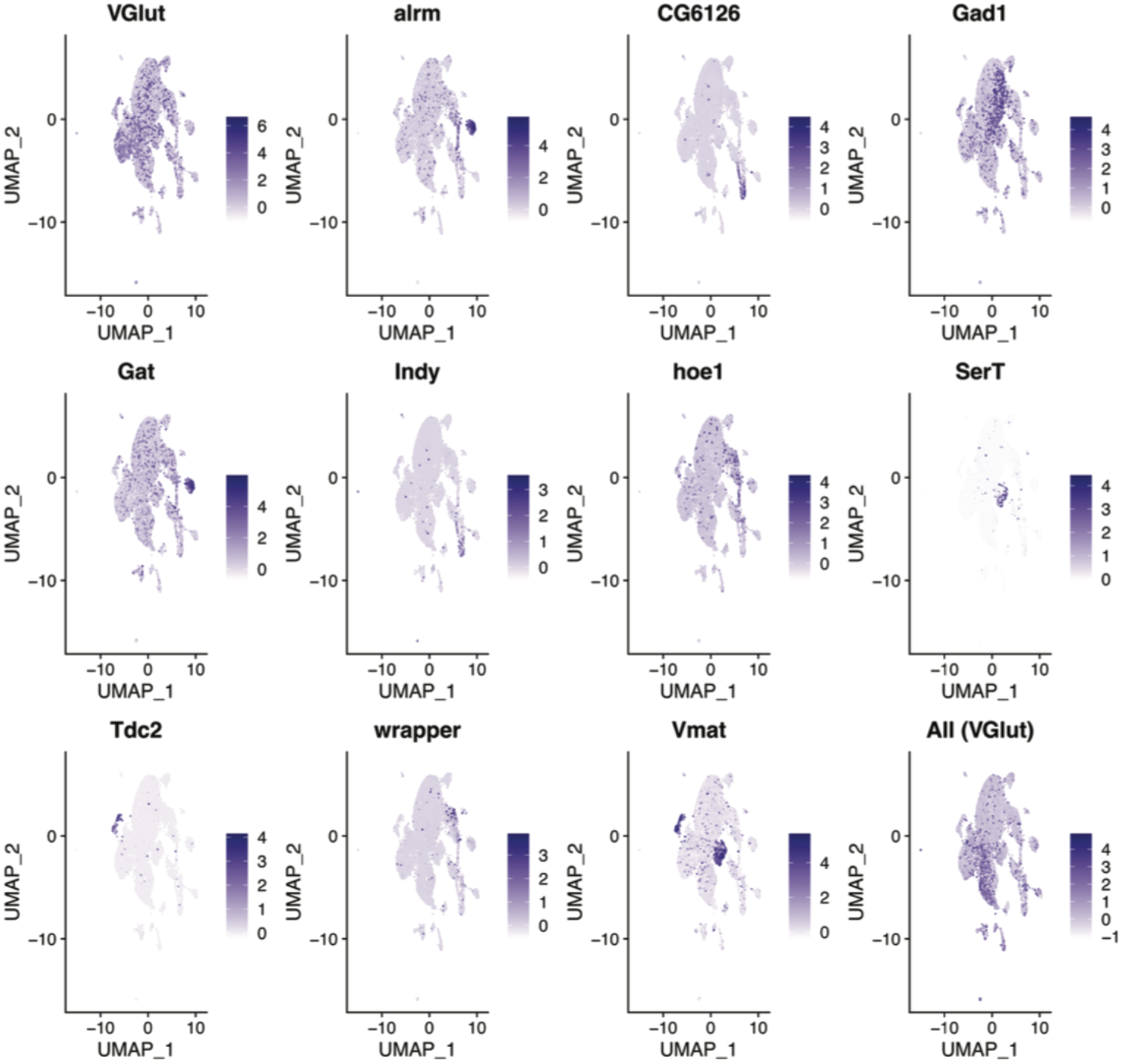
Cell-type markers in single cell RNA-seq data. Expression of different genes used as markers to define neuronal and glial populations in the larval nervous system highlighted in Figure 3D.

**Supplementary Figure 4.**
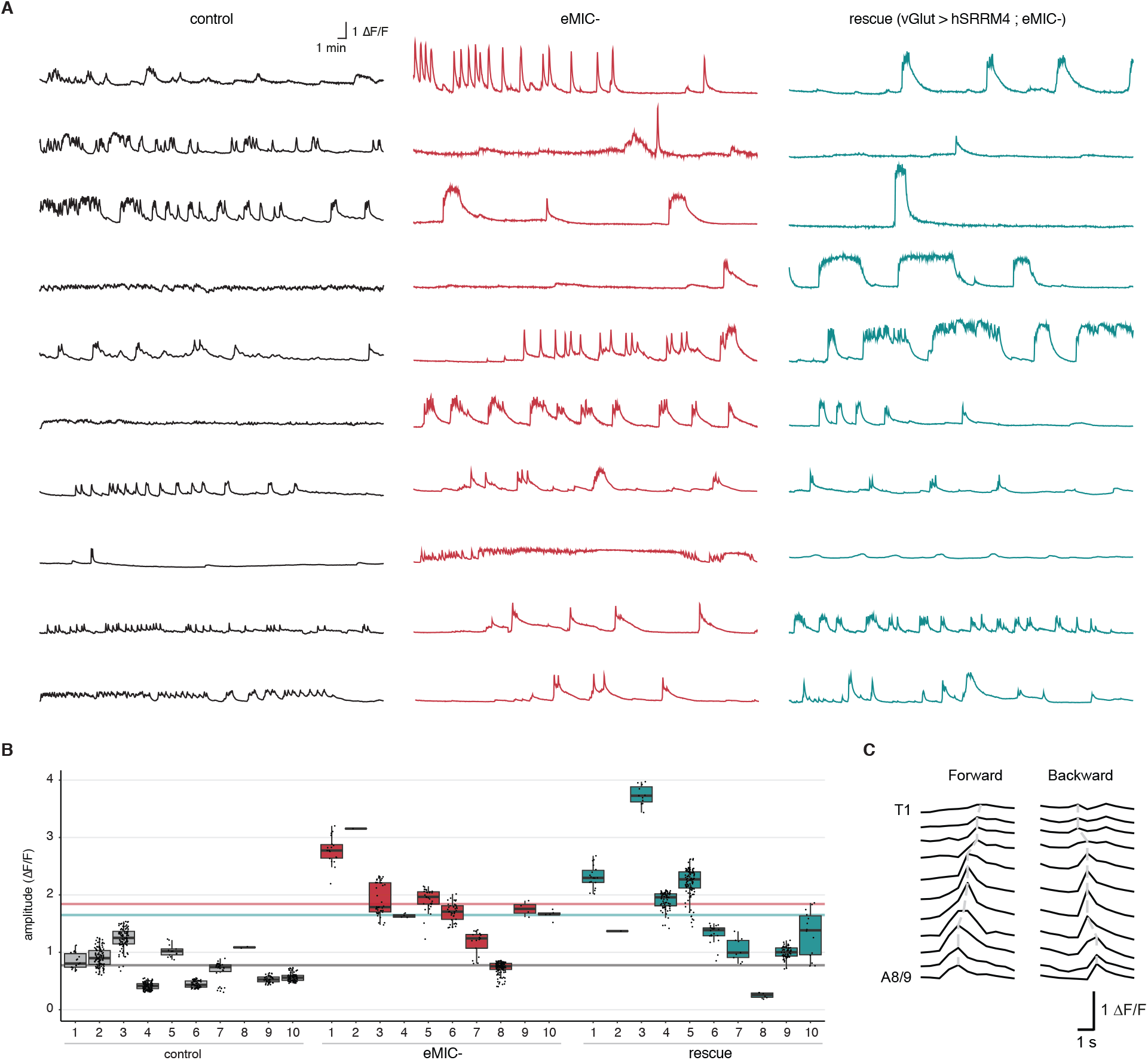
Fictive locomotion experiments in L3 larvae. **A.** Traces representing the mean activity across all segments of the ventral nerve cord (VNC) per sample. Activity is calculated based on the change in fluorescence (F) of the GCaMP7b calcium indicator over the base line (ΔF/F). **B.** Distribution of amplitudes for every peak of activity in each sample. Horizontal lines mark the average values for each genotype. **C.** Representative activity patterns across VNC segments during forward and backward waves.

**Supplementary Figure 5.**
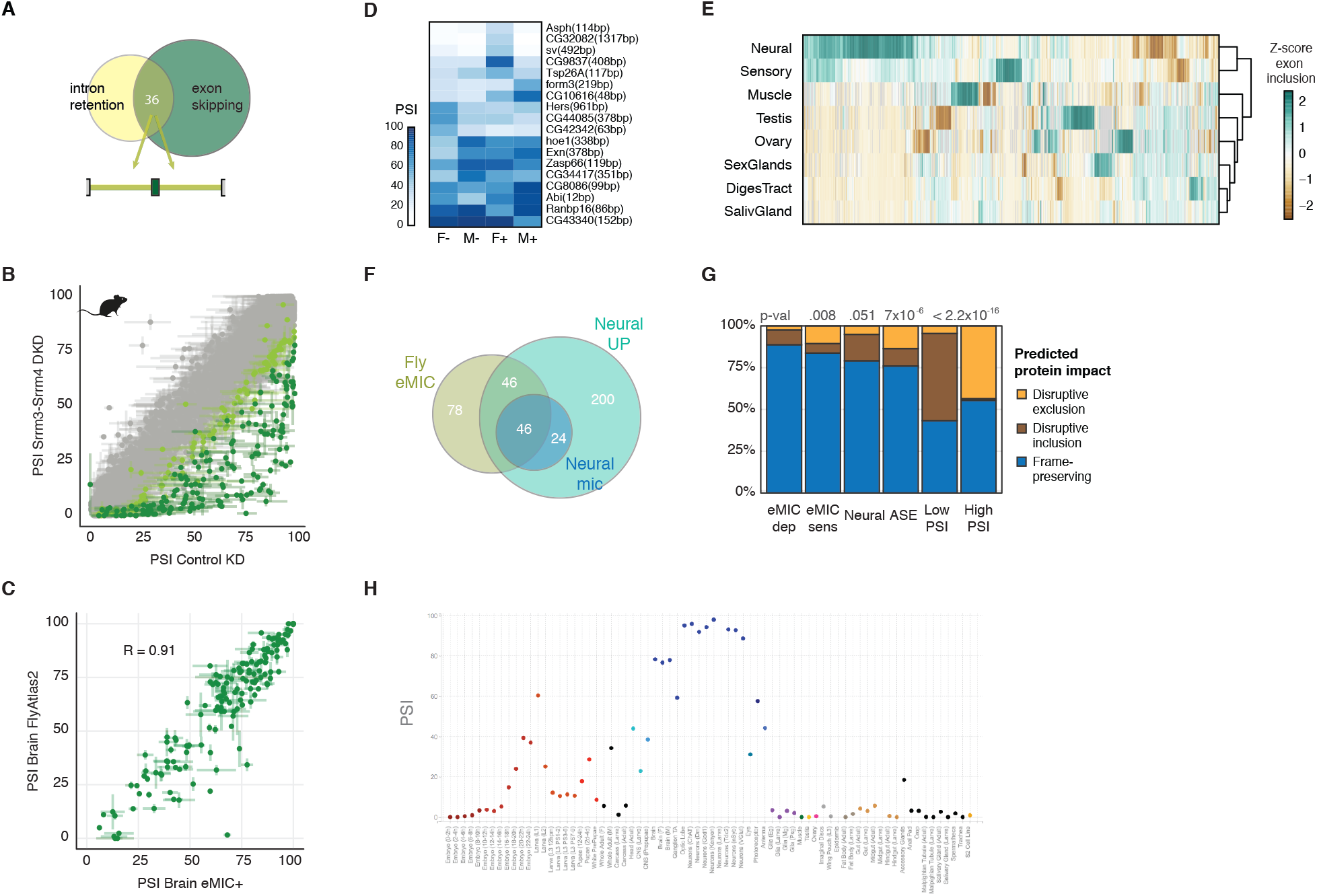
Regulation of AS by the eMIC domain. **A.** Overlap of alternatively retained introns and alternatively spliced exons (introns adjacent to identified alternatively spliced exons). **B.** PSI of mouse eMIC-dependent exons upon perturbation of Srrm3 and Srrm4 mRNA levels. Data from ^21,27^. KD: knock-down, DKD: double knock-down. **C.** PSI values of eMIC-dependent exons in our controls (x-axis) and wild-type brain samples from FlyAtlas 2 ^28^ (y-axis). Error bar ends indicate the difference between male and female samples. **D.** Inclusion levels of exons alternatively spliced between sexes in eMIC- and control adult brain samples as quantified by RNA-seq. F: female, M: male, +: eMIC+, -: eMIC-. In grey: insufficient read coverage. **E.** Tissue-specific alternative exons in *Drosophila*. Data sources included in Table S1. **F.** Overlap of eMIC-dependent exons and neural-enriched exons. Within neural exons, microexons (mic, exons shorter than 28 nt) are highlighted in dark-blue. **G.** Prediction of the effect of AS exons on their cognate proteins. dep: dependent, sens: sensitive, ASE: other AS exons, PSI: percent spliced in, Low PSI: cryptic exons, High PSI: constitutive exons. **H.** Representative example of the PSI quantification across tissues for AS events available at vastdb.crg.eu.

**Supplementary Figure 6.**
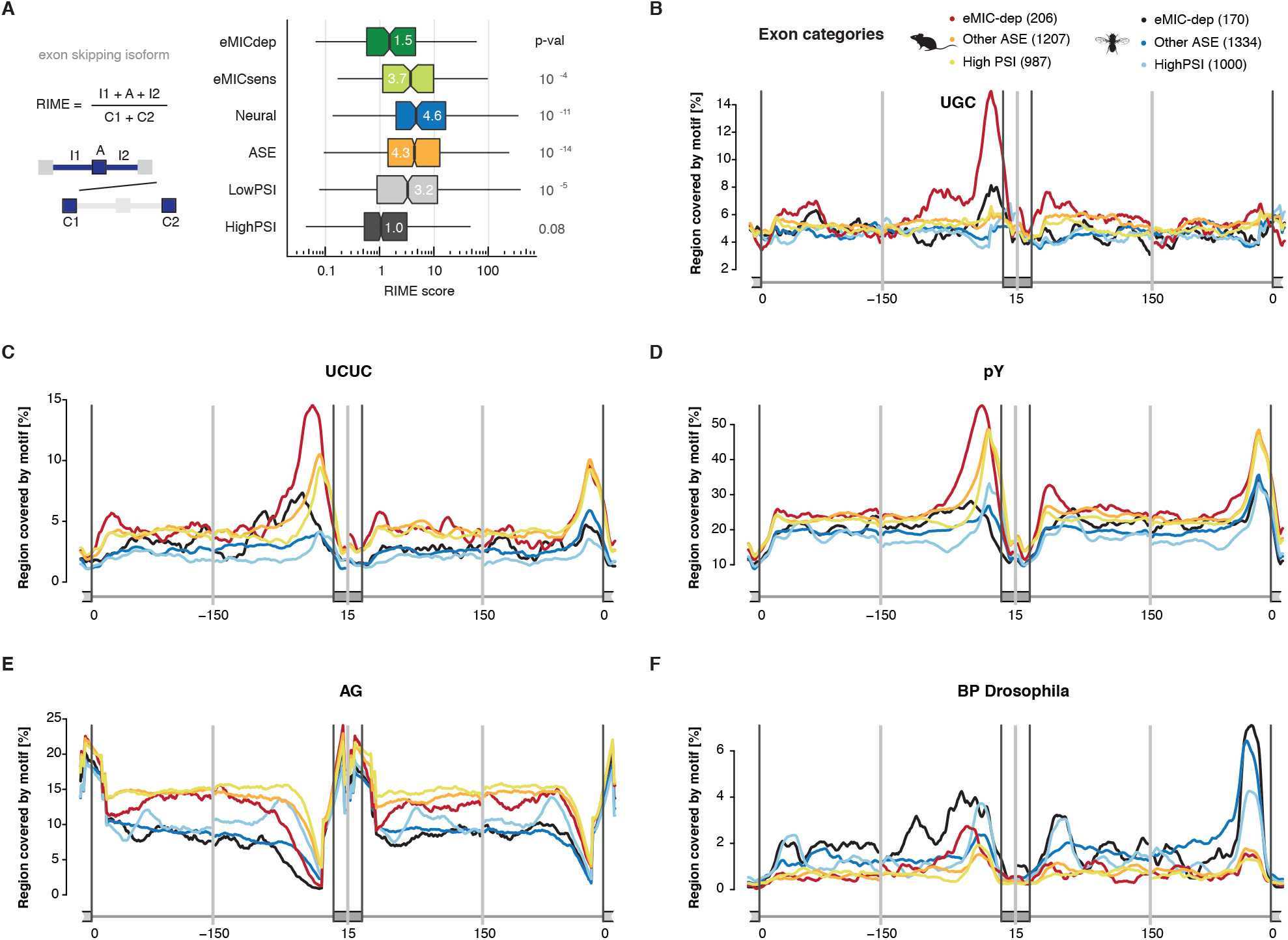
*cis*-regulatory features associated with eMIC-dependent splicing in mice and flies. **A.** Ratio of Intron to Mean Exon length (RIME) score for introns harbouring different types of cassette exons (Table S2). eMICdep and eMICsens: eMIC-dependent and eMIC-sensitive exons, ASE: other alternatively spliced exons, LowPSI and HighPSI: cryptic and constitutive exons, respectively. **B-F.** RNA-maps of motifs enriched or depleted in the intronic regions flanking eMIC-dependent exons: UGC motifs (B), alternating CU-rich tetramers (C), polypyrimidine tetramers (D), AG dinucleotide (E) and the consensus branch-point sequence in *Drosophila* CUAA[C/U] (F). Length of sliding window: 27 nt.

**Supplementary Figure 7.**
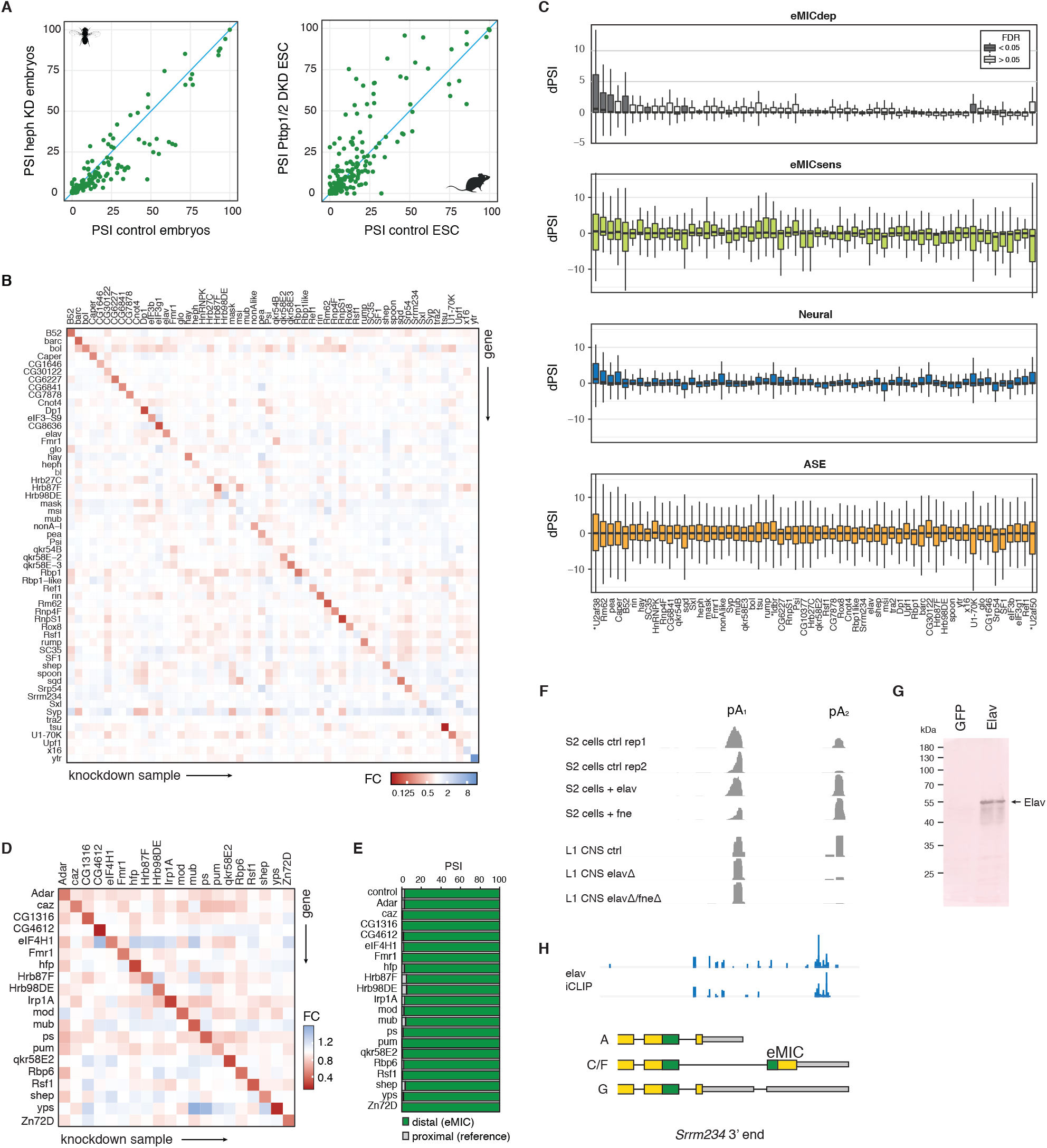
Cross-talk between eMIC splicing and other RBPs. **A.** Inclusion of eMIC-dependent exons in PSI (percent spliced in) upon perturbation of heph (Ptbp1/2/3 *Drosophila* ortholog) expression levels. Left, heph knockdown (KD) and control *Drosophila* embryos from ^39^. Right, Ptbp1/2 double knockdown (DKD) and control mouse embryonic stem cells (ESCs) from ^96^. **B,C.** Expression of RNA binding proteins (C) and change in inclusion levels (dPSI) for different types of exons upon knockdown of an array of RBPs in *Drosophila* SL2 cells (D). Data from modENCODE ^41^ or ^42^ (the latter are marked with an asterisk). FC: fold-change relative to control KD samples, eMICdep: eMIC-dependent exons, eMICsens: eMIC-sensitive exons, ASE: other alternatively spliced exons. In boxplots, centre of the box marks median values, box limits mark interquartile ranges (IQR) and whiskers 1.5 IQR. **D,E**. Expression of RBPs (E) and AS at the 3′ end of *Srrm234* (E) upon KD of several RBPs in *Drosophila* adult brains. Data from ^47^. **F.** 3′ seq reads at the *Srrm234* locus upon perturbation of *elav/fne* levels: overexpression in SL2 cells or knockout in L1 larva central neural system (CNS). pA: poly-adenylation site, ctrl: control, rep: replicate. Data from ^48^. **G.** Elav expression in SL2 cells detected by western blot. Ponceau staining was used as loading control. **H.** Elav iCLIP tags at the *Srrm234* 3’end region from adult heads. Data from ^49^. Bottom, annotated *Srrm234* isoforms based on AS and poly-adenylation at this region.

**Supplementary Figure 8.**
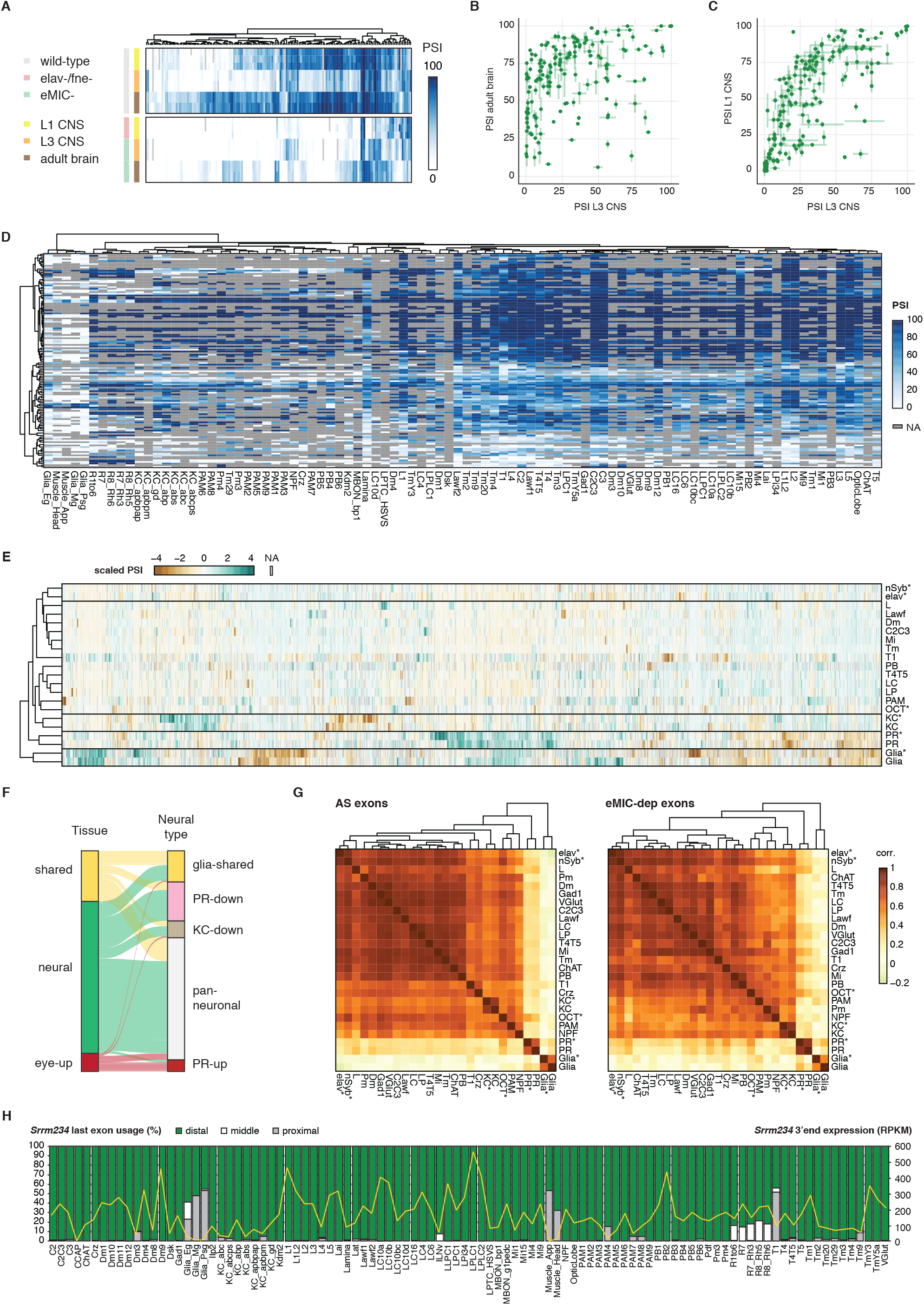
Inclusion of eMIC-dependent exons across neural cell types. **A.** Heatmap for the inclusion levels of eMIC-dependent exons in L1 and L3 CNS and adult brains for control and mutant flies. PSI: Percent Spliced In. **B.** Comparison of the PSI values of eMIC-dependent exons in larval central neural system (CNS) and adult brains. Error bar ends indicate the difference between male and female samples. Data generated in this study. **C.** PSI values of eMIC exons in L1 and L3 larva central neural system (CNS). L1 data from ^48^, error bars representing the PSI range across replicates. L3 data, from FlyAtlas 2 and this study, error bar ends indicating PSI values in each source. **D.** eMIC-dependent exon inclusion across cell types in the *Drosophila* adult optic lobe. Data from ^55^. PSI: percent spliced in. NC: no sufficient read coverage. **F.** Alluvial plot depicting the overlap between groups of eMIC-dependent exons based on their inclusion profile across tissues (left) or neural cell types (right). **E.** Inclusion levels (scaled for each exon) of all alternatively spliced (AS) exons among different cell types in the fly optic lobe. Sample sources are listed in Table S1. **G.** Sample to sample correlation (corr.) distance matrix for different cell types in the fly optic lobe using either all AS exons (left) or eMIC-dependent exons only (right). PR: photoreceptors, KC: Kenyon cells, OCT: octopaminergic neurons, nSyb: all neurons. **H.** Alternative last exon usage and gene expression levels of Srrm234 (grey line) across cell types in the optic lobe. Data from ^55^. RPKM: corrected reads per kilobase per million mapped reads.

**Supplementary Figure 9.**
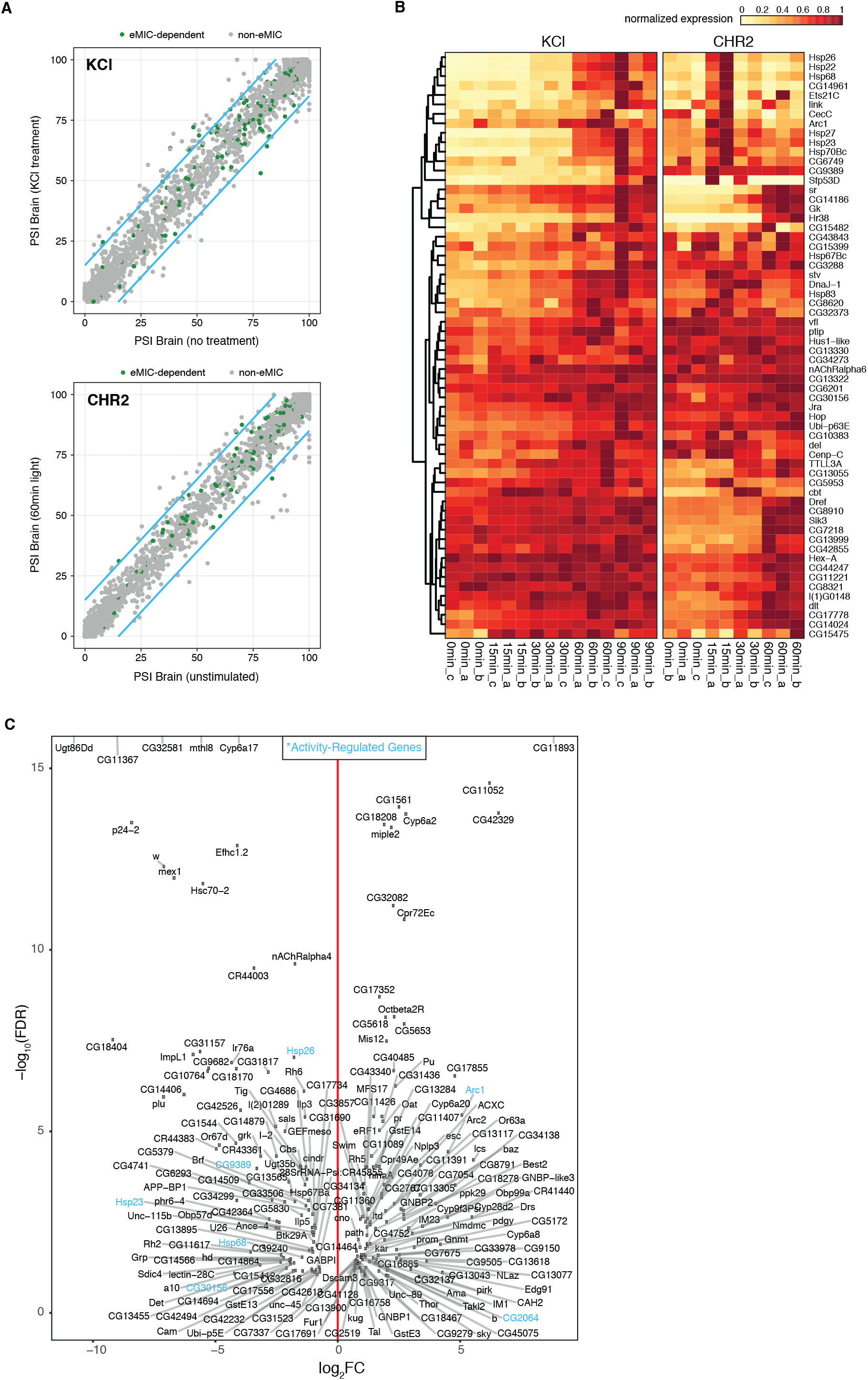
Neuronal-activity regulated transcriptomic changes and differential gene expression in eMIC-brains. **A.** PSI of genome-wide *vast-tools* exons upon two stimulation paradigms: KCl-induced depolarization (top), and pan-neuronal optogenetic activation (CHR2, bottom). Data from ^59^. **B.** Normalized expression of up-regulated genes upon two types of neuronal stimulation: KCl-induced depolarization (left) or optogenetically with CHR2 (right). **C.** Differentially expressed genes (DEG) between eMIC- and control adult brains. FC: fold-change, FDR: false discovery rate (adjusted p-values). In blue, genes regulated by sustained neuronal depolarization as identified in panel B.

**Supplementary Figure 10.**
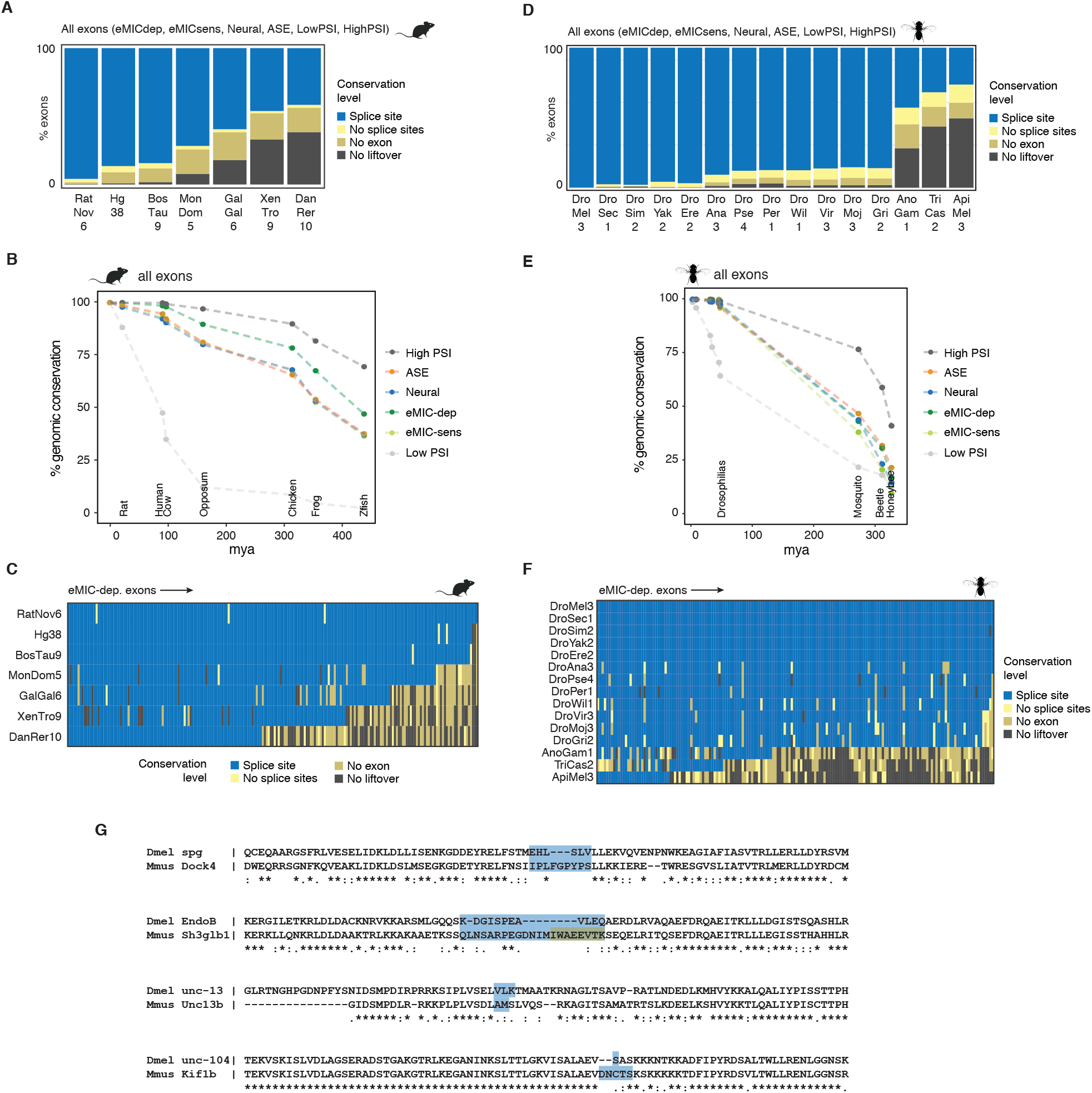
Genomic conservation of eMIC-dependent exons. **A.** Percentage of mouse AS exons conserved in the genome assemblies of other species as identified by liftOver (see Methods). **B.** Genomic conservation of different types of exons. Y-axis represents percentage of exons with identified spliced sites between the focal species (mouse) and each other species, distributed on the x-axis according to the distance to their last common ancestor with the focal species. Related to Figure 7A, but without requiring liftOver of the adjacent constitutive exons. HighPSI: constitutive exons, ASE: alternatively spliced exons, Neural: neural-enriched exons, eMIC-dep and eMIC-sens: eMIC-dependent and sensitive exons, LowPSI: cryptic exons, mya: million years ago. **C.** Genomic conservation of each mouse eMIC-dependent exon in other vertebrate species. **D-F.** Equivalent analyses to panels A-C, done using *D. melanogaster* AS exons (dm6 assembly). **G.** Protein alignment in the region surrounding the exons regulated by the eMIC domain (highlighted with coloured boxes) for the only 4 exons shared between the fly and mammalian programs. Dmel: *Drosophila melanogaster*, Mmus: *Mus musculus*.

**Supplementary Figure 11.**
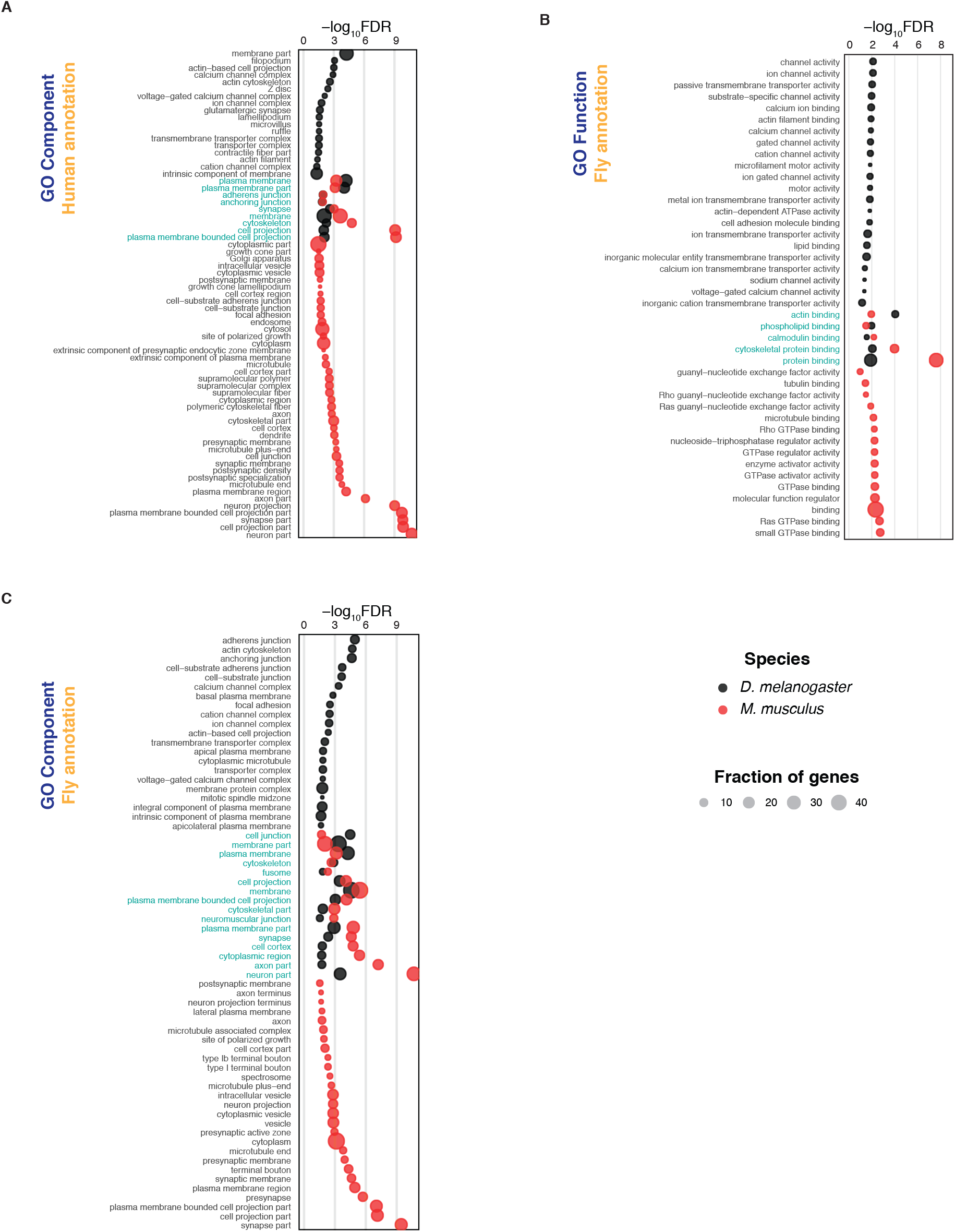
Gene ontology (GO) terms enriched in eMIC targets. **A.** GO terms in the Component categories for the human orthologs of *Drosophila* and mouse genes harbouring eMIC-dependent exons. **B,C.** Enrichment of GO terms for genes with eMIC exons, using the *Drosophila* GO annotation using either *Drosophila* genes or the *Drosophila* orthologs of mouse genes, for the Function (B) and Component (C) categories. FDR: False Discovery Rate, Fraction: percentage of genes with eMIC exons associated with each GO term. In blue, GO terms enriched in both *Drosophila* and mouse eMIC target genes.

